# Transposable Element activation promotes neurodegeneration in a *Drosophila* model of Huntington’s disease

**DOI:** 10.1101/2020.11.19.389718

**Authors:** Assunta Maria Casale, Francesco Liguori, Federico Ansaloni, Ugo Cappucci, Sara Finaurini, Giovanni Spirito, Francesca Persichetti, Remo Sanges, Stefano Gustincich, Lucia Piacentini

## Abstract

Huntington’s disease (HD) is a late-onset, autosomal dominant disorder characterized by progressive motor dysfunction, cognitive decline and psychiatric disturbances. The most prominent pathological manifestation is a selective loss of medium-sized spiny neurons of the striatum. The disease is caused by a CAG repeat expansion in the *IT15* gene, which elongates a stretch of polyglutamine at the amino-terminal of the HD protein, Huntingtin (Htt). Despite the accumulation of an impressive amount of data on the molecular basis of neurodegeneration, no therapeutic treatments are available and new pharmacological targets are needed.

Transposable Elements (TEs) are mobile genetic elements that constitute a large fraction of eukaryotic genomes. Retrotransposons (RTEs) replicate through an RNA intermediate and represent approximately 40% and 30% of the human and *Drosophila* genomes. Mounting evidence suggests that mammalian RTEs are normally active during neurogenesis and may be involved in diseases of the nervous system.

Here we show that TE expression and mobilization are increased in a *Drosophila melanogaster* HD model. By inhibiting TE mobilization with Reverse Transcriptase inhibitors, polyQ-dependent eye neurodegeneration and genome instability in larval brains are rescued and fly lifespan is increased. These results suggest that TE activation may be involved in polyQ-induced neurotoxicity and a potential pharmacological target.

## Introduction

Huntington’s disease (HD) (OMIM #143100) is a late-onset, autosomal dominant disorder characterized by progressive motor dysfunction, cognitive decline and psychiatric disturbances, which lead to death approximately 15-20 years after the initial symptoms (Folstein 1989). The most prominent pathological manifestation of the disease is a selective and gradual loss of medium-sized spiny neurons of the striatum, although, in the late stage of the disease, pathological alterations occur in additional brain regions (Vonsattel and DiFiglia 1998). The disease is caused by a CAG repeat expansion in the *IT15* gene, which elongates a stretch of polyglutamine (polyQ) at the amino-terminal end of the HD protein, Huntingtin (Htt) (HDCRG 1993). The number of CAG repeats varies between 6 and 35 units on normal chromosomes, whereas on HD chromosomes the repeat is expanded above the pathological threshold of 36 CAGs and can range as high as 150 or more. Htt is ubiquitously expressed and it is mainly, but not exclusively, localized to the cytoplasm where it may associate to organelles (DiFiglia et al. 1995). Htt interacts with numerous proteins implicated in processes as diverse as gene transcription, RNA splicing, intracellular transport, signal transduction and metabolism.

The HD mutation, by virtue of the expanded glutamine tract, confers a novel toxic property to the mutant protein. A gain of function mechanism is indeed supported by genetic and experimental data (Duyao et al. 1995; Nasir et al. 1995). In HD target neurons, the trigger event driven by mutant Huntingtin is likely to occur many years before the appearance of the first signs of neurodegeneration. The identification of genetic modifiers of the age of onset has established the role of somatic mosaicism of the CAG trinucleotide repeats in neural genomes (Gusella et al. 2014; Lee et al. 2019). Mutant Huntingtin triggers extensive epigenetic-chromatin deregulation as shown by the analysis of DNA methylation and histone modifications (Seong et al. 2010; Bassi et al. 2017). The disease cascade that leads to neuronal cells’ death involves several pathways including defects in the proteasome apparatus, glutamate-mediated excitotoxicity, mitochondrial dysfunction and neuroinflammation (Ross and Tabrizi 2011).

Our understanding of the molecular basis of HD pathogenesis remains incomplete and a treatment for this devastating disease is still not available. It is therefore important to identify previously unnoticed pathways that may be altered in HD and potential target of therapeutic treatments.

*Drosophila melanogaster* is an excellent choice for modelling HD pathology and disease mechanisms. It has a relatively simple central nervous system with an architecture that separates specialized functions such as vision, olfaction, learning and memory similarly to mammalian nervous systems (Armstrong et al. 1995). Moreover, the majority of pathological features of HD can be recapitulated in transgenic fly models, including dominant gain of function neurotoxicity, nuclear inclusion formation, progressive neurodegeneration, behavioural abnormalities and early death (Chan et al. 2002).

In the last few years, correlations between transposon activation and neurological diseases have been observed (Tam et al. 2019; McConnell et al. 2017). Transposable Elements (TEs) are mobile genetic elements that constitute a large fraction of eukaryotic genomes (Belancio et al. 2008). Retrotransposons (RTEs) replicate through an RNA intermediate by taking advantage of reverse transcriptase (RT) activity and represent approximately 40% and 30% of the human and *Drosophila* genomes, respectively. During the co-evolution of TEs with their host genomes, organisms have evolved efficient mechanisms to prevent and regulate TE activation and mobilization including methylation and Piwi-interacting RNAs (piRNAs) expression (Ozata et al. 2019). Mounting evidence shows that Long Interspersed Nuclear Element 1 (LINE-1) sequences are normally active during neurogenesis in both rodent and human tissues and somatic mobilization of TEs has been observed in different human brain regions giving rise to mosaicism (Bodea et al. 2018). The extent and the functional role of TE mobilization in brain physiology remain unclear (Evrony et al. 2016; Upton et al. 2015; Cappucci et al. 2018). LINE-1 mobilization has also been associated to deletion of genomic fragments in human brains (Rodriguez-Martin et al. 2020; Erwin et al. 2016). Emerging evidence suggests an association between unregulated activation of TEs and diseases of the nervous system (Jacob-Hirsch et al. 2018). Pathological TE activation has been observed in, among others, Rett syndrome (Muotri et al. 2010), ataxia telangiectasia (Coufal et al. 2011), amyotrophic lateral sclerosis (ALS) (Li et al. 2012; Krug et al. 2017), Alzheimer’s disease (AD) (Guo et al. 2018; Sun et al. 2018) and Parkinson’s disease (Blaudin de Thé et al. 2018).

In a *Drosophila* model of ALS with the ectopic expression of human TAR DNA-binding protein 43 (TDP-43), a de-repression of both LINE and LTR families of TEs was observed (Krug et al. 2017; Li et al. 2012). The degenerative fly phenotypes were rescued by pharmacologically inhibiting RT activity or by interfering with the expression of the endogenous retrovirus (ERV) *gypsy* (Krug et al. 2017). *Gyspy*, together with *copia* and *Het-A* RNAs, was also found induced in a fly model of AD over-expressing wild-type or mutant Tau (Guo et al. 2018). Interestingly, chromatin decondensation and decreased expression of piwi/piRNAs led to TE activation in another fly model of tauopathy where inhibition of RT activity was able to rescue the neurodegenerative phenotype (Sun et al. 2018).

These data suggest that resurrection of TEs could be causally involved in neurodegeneration and recapitulated in fly models of disease.

By taking advantage of a well-established fly HD model, here we show that TE activation may also be involved in HD pathogenesis and its inhibition may represent a new therapeutic strategy.

## Results

### TE expression is induced in HD fly model

In this study we take advantage of the *Drosophila* HD experimental model generated by Romero and collaborators (Romero et al. 2008). It is based on the expression of full-length human Huntingtin (Htt) protein with 128 glutamines (*128QHtt^FL^*, pathogenic HD construct) under the control of the UAS-Gal4 system (Brand and Perrimon 1993). In this bipartite system, the expression of the HD pathogenic construct placed downstream of UAS sequences depends on the spatial and temporal expression of the yeast transcriptional activator Gal4. In order to induce a pan-neuronal expression of the pathogenic HD construct, we crossed flies carrying the *UAS-128QHtt^FL^* transgene (hereafter referred to as 128QHtt) to flies *elav-Gal4* driving the expression of the pathogenic construct in every post mitotic neuronal cell, starting from very early stages of development (Robinow and White 1988). We first checked by semi-quantitative RT-PCR the correct expression of *128QHtt* mRNA in *Drosophila* HD heads (Fig. 1A), we then analysed different transposon transcripts in larval and adult brain comparing transposon RNA levels from HD flies (*elav-Gal4>128QHtt*) to those from controls (*elav-Gal4/+*) (Fig. 1B). The time points for this analysis (0-2 and 10-12 days) were chosen taking into account the dynamic of survival curves indicating that HD flies have a dramatically reduced lifespan and begin to die shortly after 10 days of age (Supplemental Fig. S1A).

**Figure 1.**
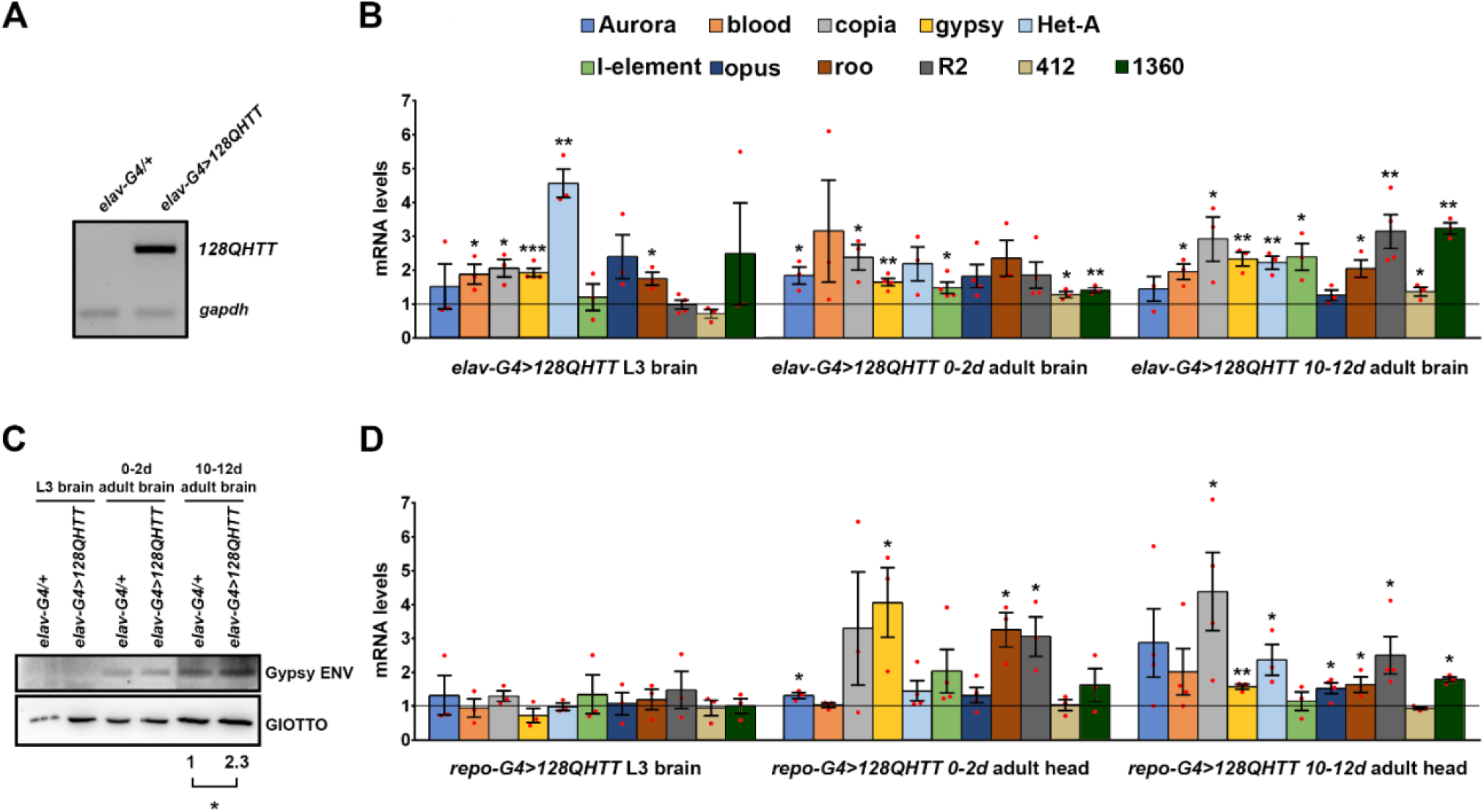
Neuronal and glial expression of 128QHtt results in transposable element de-repression. (A) Semi quantitative RT-PCR analysis to assess the induction levels of the HD transgenic construct by elav-Gal4. cDNA was prepared from total RNA purified from HD (*elav-G4>128QHtt*) and control (*elav-G4/+*) head tissues. The constitutive *gapdh* was examined as an endogenous control. (B) qRT-PCR analysis of transposable element expression in larval and adult brains of flies expressing 128QHtt in neurons (*elav-G4>128QHtt*); adult brains were analysed at both young (0-2 days) and aged (10-12 days) time points; transcript levels were normalized to *rp49* and displayed as fold change relative to flies carrying the elav-Gal4 driver with no 128QHtt transgene (*elav-G4/+*). (C) Western blot assay of Gypsy Envelope protein (ENV) expression in HD larval and adult brains. GIOTTO protein was used as a loading control. Result was expressed as means for at least three independent biological replicates (*p < 0.05, One-sample t-test). (D) qRT-PCR analysis of TE expression in larval brains and adult heads isolated from 0-2 and 10-12-days-old flies expressing 128QHtt with the pan-glial repo-Gal4 driver *(repo-Gal4>128QHtt).* Transcript levels were normalized to *gapdh* and displayed as fold change relative to flies carrying the repo-Gal4 driver with no128QHtt transgene (*repo-G4/+*). (B,D) Bar graph represents the mean ± SEM from at least three independent experiments (*p < 0.05; **p < 0.01; ***p < 0.001, Unpaired t-tests). Red dots indicate individual data points. The black horizontal line indicates the Fold Change control value, set to 1.

We found that different classes of TE transcripts were significantly upregulated in HD neurons at both larval and adult stages (Fig. 1B). *Het-A* was the most induced class at the larval stage while several classes presented similar induction at 0-2 and 10-12 days of adult brain. Interestingly, the increase of TE transcripts at 10-12 days correlated with age-dependent neurodegeneration since at this stage an extensive neuronal loss was evident, as demonstrated by the strong decrease of the neuronal marker Elav (Supplemental Fig. S1B). In addition, consistently with the expression of *Gypsy* transcripts, a significant age-dependent increase of the Gypsy ENV (envelope) protein levels was observed in adult brains expressing pan-neuronally 128QHtt (Fig. 1C). In order to further confirm the TE activation was actually due to expression of the polyQ-expanded 128QHtt, we also examined the effect on TE deregulation of a wild-type full-length Htt containing 16 glutamines (16QHtt). As shown in the Supplemental Fig. S2, the absence of significant differences between *elav-Gal4/+* and *elav-Gal4>16QHtt* demonstrates that TE activation shown in Fig. 1B is due to expression of the pathogenic polyQ-expanded 128QHtt transgene and not to the presence of a wild-type Htt transgene.

Since glial cell lineage expression of human mutant Htt (hHtt103Q) was previously shown to induce developmental and late-onset neuronal pathologies in *Drosophila* model (Tamura et al. 2009), we expressed the pathogenic HD protein in glial cells by using the pan-glial driver repo-Gal4 (Awasaki et al. 2008) and tested TE RNA induction. *Repo-Gal4>128QHtt* raised at 29 °C showed severe locomotive defects and very early death; over 70% of the *repo-Gal4>HD* flies died within 4-5 days following eclosion and the survivors died more gradually over the ensuing 8-10 days. Aiming to analyse TE expression in *repo-Gal4>128QHtt* adult heads, we raised *repo-Gal4>128QHtt* flies at 25°C to partially suppress Gal4 activity and increase their survival rate (as summarized in Supplemental Fig. S3A). Pan-glial expression of mutant Htt induced extensive glial degeneration, especially at 10-12 days post-eclosion (Supplemental Fig. S3B). By qRT-PCR we analysed RNA samples from larval brains and whole heads of *repo-Gal4>128QHtt* flies collected at both 0-2 and 10-12 days post-eclosion. As shown in Fig. 1D, while no significant increase in *repo-Gal4>128QHtt* larval brains was observed, several TE classes were upregulated *in repo-Gal4>128QHtt* adult heads with a different pattern of expression when compared to *elav-Gal4>128QHtt*.

### Loss of histone H3 trimethylation at Lys9 (H3K9me3) and global heterochromatin relaxation contribute to TE silencing release in HD

To investigate the molecular mechanisms by which HD might contribute to transposon activation, we analysed, by chromatin immunoprecipitation coupled to quantitative PCR (ChIP-qPCR), the levels of H3K9me3 on promoters and coding regions of up-regulated TEs. Trimethylation of histone H3 on lysine 9 is a specific epigenetic mark of heterochromatic domains and it is essential for heterochromatin formation and maintenance (Allshire and Madhani 2018). As shown in Fig. 2A, we found a drastic reduction of H3K9me3 occupancy in adult heads isolated from 10-12-day-old *elav-Gal4>HD* flies for almost all sequences of TE analysed, especially on their promotors.

**Figure 2.**
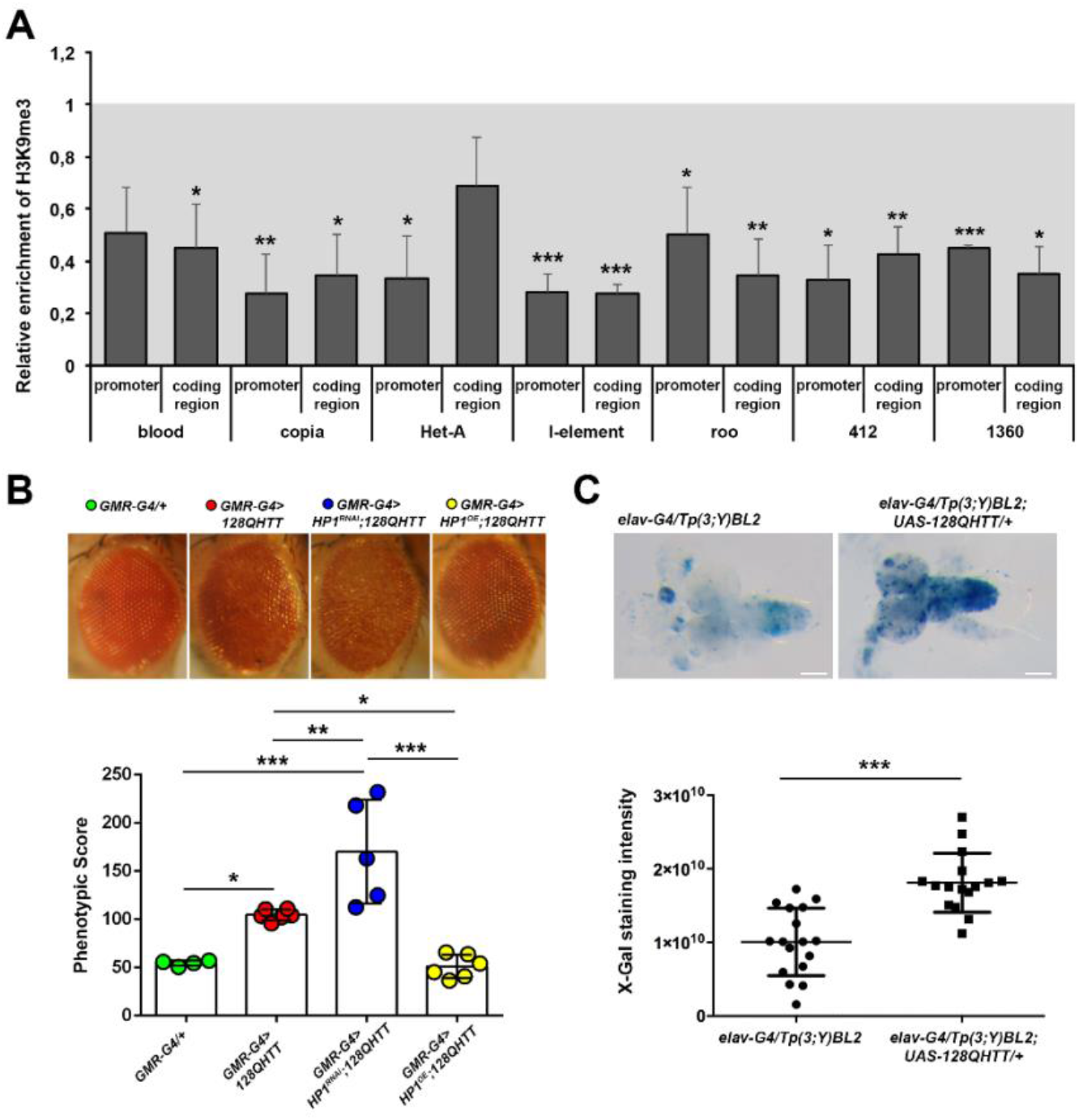
Global heterochromatin relaxation mediates HD-induced TE activation. (A) Transposon DNA sequences show decreased H3K9me3 levels in HD. The relative fold enrichment, normalized to the heterochromatic F22 control region, was calculated by dividing the amount of immunoprecipitated DNA from *elav-G4>128QHtt* heads by that from the control *(elav-G4/+).* Bars represent mean ± SEM of three independent experiments performed in duplicate (*p < 0.05; **p < 0.01; ***p < 0.01, Unpaired t-tests.) The grey area represents the control value set to 1. (B) HP1 is a genetic modifier of HD eye phenotype. dsRNA-mediated inactivation of HP1 (*GMR-Gal4>HP1^RNAi^;128QHtt*) substantially enhances depigmentation and structural alterations of HD eye while HP1 overexpression (*GMR-Gal4>HP1^OE^;128QHtt*) strongly prevents HD-induced eye neurodegeneration. *GMR-G4/+* was used as control. The lower graph represents the phenotypic scores of each indicated genotype. The phenotypic scores are concordant with the visual assessment of the eye phenotypes. The number of images used for this analysis ranged from 4 to 6 (*p < 0.05; **p < 0.01; ***p < 0.001, One-way ANOVA with Tukey’s post hoc test). (C) Mutant Htt suppresses BL2 PEV. Histochemical staining for β-galactosidase activity in larval brains isolated from HD (*elav-G4/Tp(3;Y)BL2; UAS-128QHtt*, right panel) and control (*elav-G4/Tp(3;Y)BL2*, left panel) male larvae. Scale bar indicates 100μm. Scatter plot indicates the quantitative analysis of X-gal staining intensities. Data are the means ± SD. The dots indicate individual data points (***p < 0.01, Unpaired t-tests).

The overall reduction in H3K9me3 levels on TE sequences in HD could result from a global heterochromatin decondensation, a phenomenon recently reported in *Drosophila* models of tauopathy (Sun et al. 2018; Frost et al. 2014). To verify whether constitutive heterochromatin widespread relaxation represents a distinctive feature also in the HD model, we analysed the effects of perturbing the expression of HP1, a genetic interactor of *dHtt* and a key positive regulator of heterochromatin maintenance (Eissenberg et al. 1990; Dietz et al. 2015; James and Elgin 1986).

To this purpose, we took advantage of the ability of 128QHtt to elicit photoreceptors degeneration upon its expression in the developing eye using the GMR-Gal4 driver (Romero et al. 2008; Ellis et al. 1993). The examination of the external eye morphology using bright-field microscopy revealed that overexpression of HP1 in HD genetic background induces a significant recovery of pigmentation and structure of HD eye while its functional inactivation results in worsened eye phenotypes (Fig. 2B). To quantify the degree of severity of the eye phenotypes, we used Flynotyper (Iyer et al. 2016), a computational method that calculates a phenotypic score based on the disorganization or altered symmetry of the ommatidial arrangement.

These results confirm that changes in heterochromatin structure play a central role in HD pathogenesis.

In *Drosophila*, a widely accepted genetic tool in understanding the dynamic of heterochromatin state is the Position-Effect Variegation (PEV), a heterochromatin-induced gene silencing that occurs when an euchromatic gene, juxtaposed with heterochromatin by chromosomal rearrangements or translocations, is transcriptionally silenced in some cells but active in others, producing a characteristic mosaic phenotype (Wakimoto 1998). Genes or transgenes that impact heterochromatin structure and organization act as dose-dependent modifiers of PEV-based gene silencing (Elgin and Reuter 2013). For instance, loss of *dHtt* and HP1 mutations have been previously shown to suppress PEV (Eissenberg et al. 1990; Dietz et al. 2015), leading to reduced silencing of variegating genes, whereas HP1 overexpression resulted in an increased heterochromatic gene silencing (Eissenberg et al. 1992). To assay whether 128QHtt expression can induce heterochromatin decondensation, thus suppressing PEV, we used the *Tp(3;Y)BL2*, a chromosome rearrangement carrying the *Hsp70-lacZ* inducible variegating transgene into Y pericentromeric heterochromatin. This can be used to detect PEV modifications in larval tissues upon staining for β-galactosidase activity (Lu et al. 1996). To analyse PEV in larval brain tissues, we crossed *elav-G4* females to *Tp(3;Y)BL2* or *Tp(3;Y)BL2; UAS-128Htt* males. We then compared *elav-G4/Tp(3;Y)BL2* and *elav-G4/Tp(3;Y)BL2; UAS-128Htt/+* F1 males for Lac-Z expression in larval brains (Fig. 2C). X-Gal staining revealed that the expression of 128QHtt strongly suppresses the position-effect variegation in the *Tp(3;Y)BL2* line (Fig. 2C).

In summary, evidence in larval and adult brains and in the eye strongly suggest that mutant Htt induces TE activation through a widespread heterochromatin decondensation.

### RT inhibitors rescues HD eye phenotypes and HD-induced genome instability

To assess the functional role of TE mobilization in poly-Q induced neurotoxicity, we adopted a pharmacological approach to block the transposition of RTEs with RT inhibitors and monitored their effects on the HD eye phenotype. Flies expressing *GMR-Gal4>128QHtt* were fed over the entire developmental period on standard cornmeal agar yeast medium supplemented with 1mg/mL Lamivudine (3TC) or 5mg/mL Zidovudine (AZT), two well-known RT inhibitors (Goic et al. 2013; Wood et al. 2016; Sun et al. 2018; Krug et al. 2017) and therefore blockers of RTEs transposition. As shown in Fig. 3A, 3TC and AZT treatments strongly rescued the altered HD eye phenotype in 100% of HD flies carrying only one copy of HD mutagenic construct (*GMR-Gal4/+; UAS-128QHtt/+*) (AZT, n=60; 3TC, n=55). In addition, AZT treatment resulted more effective because, unlike 3TC, was able to rescue the HD eye phenotype also in 56% of flies carrying two copies of HD mutagenic construct (*GMR-Gal4/+; UAS-128QHtt/UAS-128QHtt*) (AZT, n=64; 3TC, n=46). AZT was therefore used in all subsequent experiments.

**Figure 3.**
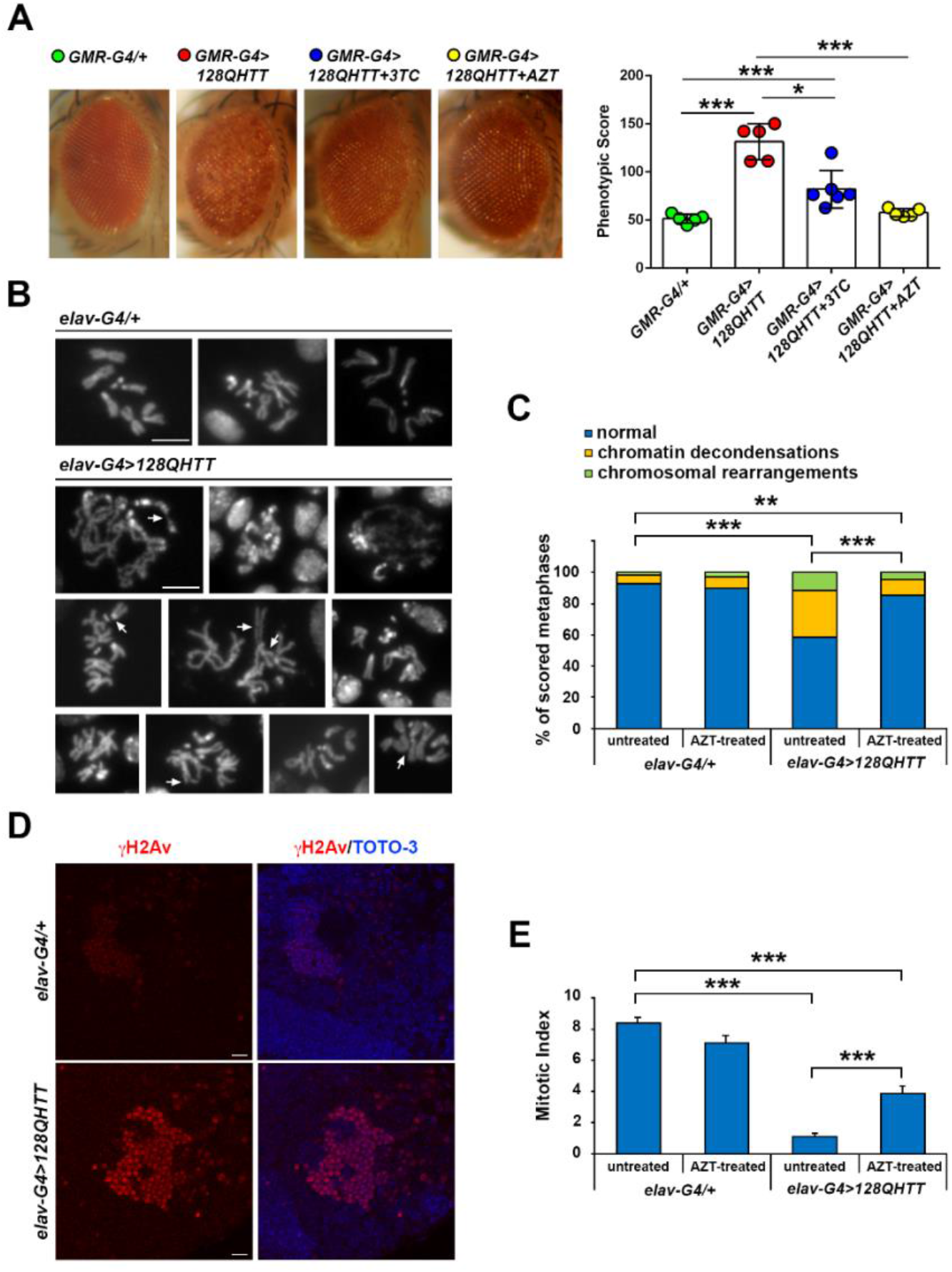
Treatment with reverse transcriptase inhibitors rescues HD eye phenotype and prevents HD-induced genome instability. (A) 3TC and AZT treatments significantly improve the altered HD eye-phenotype. Transgenic flies expressing *GMR-Gal4>128QHtt* and treated with 3TC at 1mg/mL or AZT at 5mg/mL for the entire development period, show a significant improvement of the polyQ-induced neurodegenerative phenotype relative to untreated flies. *GMR-G4/+* was used as control. The right graph indicates the phenotypic scores of each genotype. The phenotypic scores are concordant with the visual assessment of the eye phenotypes. The number of images used for this analysis ranged from 5 to 6 (*p < 0.05; **p < 0.01; ***p < 0.001, One-way ANOVA with Tukey’s post hoc test). (B) Mitotic chromosomes from third instar larval HD brains stained with DAPI. HD chromosomes show a higher degree of chromatin decondensation and structural rearrangements (see arrows for examples) when compared to the control. Scale bar indicates 5μm. (C) Quantification of chromosomal abnormalities observed in HD larval brains. (**p<0.01, ***p<0.001, Chi-square test). (D) Confocal microscopy images showing immunofluorescence against γH2Av on HD (*elav-Gal4>128QHtt*) and control (*elav-Gal4/+*) larval brains. Images were captured at 63X magnification. Scale bar indicates 10μm. (E) Bar graph represents Mitotic Index (percentage of cells in mitosis per optical field, at 40X magnification) observed in the brains of HD and control larvae treated or not with AZT (***p<0.001, One-way ANOVA with Tukey’s post hoc test).

PolyQ-expansion mutations in HD have been recently shown to lead to chromosomal instability with a molecular mechanism that remains unclear (Ruzo et al. 2018). It is well known that transposition events could represent a potential cause of genomic instability (Ayarpadikannan and Kim 2014; Belgnaoui et al. 2006; Gasior et al. 2006). Therefore, in order to determine whether HD-induced TE activation can impact genomic stability, we analysed metaphase chromosomes obtained from *elav-Gal4>128QHtt* larval brains and control ones (*elav-Gal4/+*), showing that HD metaphases present abnormal chromosome configurations such as breakages, fusions, chromosomal rearrangements and a higher degree of chromatin decondensation (Fig. 3B). Interestingly, the frequencies of chromosome abnormalities observed in HD larval brain, were significantly decreased after AZT pharmacological treatment (Fig. 3C) proving that TE activation participates in HD-induced genomic instability, as previously shown for TE in other biological contexts (Ayarpadikannan and Kim 2014).

As a confirmation of the presence of genomic damage, immunofluorescence analysis was performed on HD larval brains using an antibody against the histone variant γH2Av, an early and specific marker of DNA damage. Confocal microscopy images clearly showed that HD larval brains accumulate γH2Av-positive foci, confirming HD-induced genomic instability (Fig. 3D). This was consistent with a drastic reduction in mitotic index, indicating that DNA damage may also affect mitotic entry or, more in general, cell cycle progression. Notably, as reported in Fig. 3E, the mitotic index in AZT-treated HD larvae results significantly higher than in untreated HD larvae. All together, these results confirm a functional role of RTEs mobilization in HD pathogenesis.

### RT inhibitors extend lifespan of HD flies

In order to study a systemic effect of AZT inhibitor, we performed lifespan assays for *elav-Gal4>128QHtt* flies treated (or not) with AZT. We crossed parental lines on normal medium and transferred the HD offspring on AZT supplemented medium, starting the treatment at 0-2 or 4-5 days after eclosion (Fig. 4A,B). The log-rank analysis of lifespan curves showed that starting the AZT treatment at 0-2 days after eclosion significantly increased the median lifespan by 15.36% (the median lifespan was 15.43 ± 0.31 days for untreated flies vs 17.80 ± 0.38 days for AZT treated flies) (Fig. 4A). On the contrary, starting the AZT treatment at 4-5 days after eclosion did not ameliorate the shortened lifespan of HD flies (the median lifespan was 16.53 ±0.28 days for untreated flies vs 16.27 ± 0.32 days for AZT treated flies) (Fig. 4B). These results strongly suggest that AZT treatment is effective only starting from early stages of HD pathogenesis, while has no effect at late stages of the disease. To extend these results, we crossed the parental lines directly on AZT-medium letting the HD offspring develop on AZT-supplemented medium until post-feeding larval stage, when larvae stop feeding and commit to metamorphosis. After eclosion, HD flies were transferred to vials containing normal food medium and the survivorship was determined. Also in this case, we find a significant rescue effect in median lifespan (+7,8%) between treated and untreated HD flies (the median lifespan was 22 ± 0.39 days for untreated flies vs 23.72 ± 0.37 days for AZT treated flies) (Fig. 4C) suggesting that AZT treatment was more effective in rescuing median lifespan only when it is administered in larvae and early adult stage.

**Figure 4.**
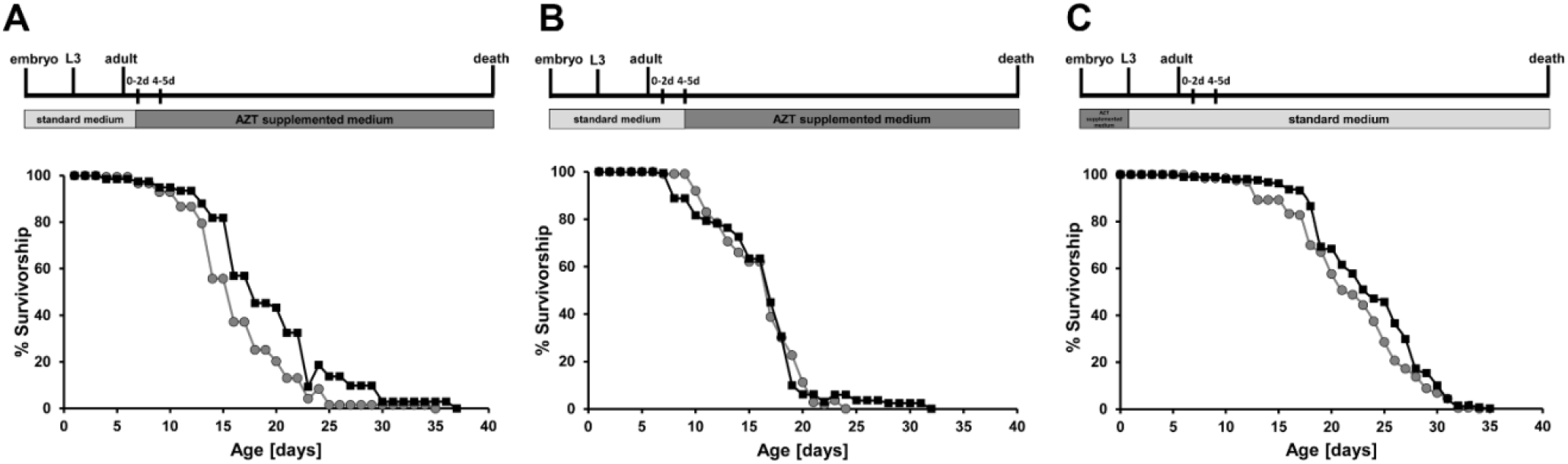
AZT treatment is more effective in rescuing median lifespan only when it is administered in larval and early adult stage. Survival curves of *elav-Gal4>128QHtt* flies untreated (grey circles) or treated with AZT (black squares) starting at 0-2 (A) or 4-5 days (B) following eclosion. The survival curves shown in (A) result significantly different (***p <0.001, Log rank-test). In (B) the survival difference is not statistically significant (p=0.68). In (C), survival curves of *elav-Gal4>128QHtt* flies untreated (grey circles) or treated with AZT (black squares) from eggs hatch to post feeding larval stage. The log rank test showed that the two curves were significantly different (**p < 0.01). The schemes at the top represent an outline of experimental strategy. Lifespan experiment was conducted with at least 150-200 flies per condition.

### WGS analysis of Structural Variants (SVs) in brain genomes of HD and control samples

To better investigate the role played by TEs in the genome instability of HD fly models, we performed whole genome sequencing (WGS) on one μg of DNA from offspring control (*elav-Gal4/+*, offspr. CTR) and HD brain tissues (*elav-Gal4>128QHtt*, offspr. HD) at 3 different developmental time points (larvae, young and aged adults). Given the size of the *Drosophila melanogaster* genome, about 60 individuals were sequenced. The 6 DNA samples were sequenced using paired-end (PE) Illumina technology. Raw reads were assessed for quality by using FastQC (Andrews 2010) and the 125bp PE reads were mapped to the *Drosophila* genome (UCSC dm6 version (Casper et al. 2017)) using *bwa mem* (Li and Durbin 2009) with default parameters. On average, >90% of the reads of the 6 samples mapped properly. The coverage calculation highlighted an average value of about 50X. Having assessed the good quality of the reads and the mappings, we took advantage of Lumpy (Layer et al. 2014) to detect SVs including break ends (BND) (indicating complex rearrangements), deletions (DEL), duplications (DUP) and inversions (INV) and of Freebayes (Garrison and Marth 2012) to detect SNP and indels (SNP). Furthermore, we carried out a TE non-reference analysis of *de-novo* Insertional Sites (ISs) by the Mobile Element Locator Tool (MELT) (Gardner et al. 2017). After normalization of the raw number of SVs, SNP-indels and TE ISs on the total number of sequenced reads, results showed a higher number of all variation types in offspring CTR with respect to HD (Supplemental Fig. S4). However, when their numbers were quantified on each chromosome, we found that chr3 (chr3L and/or chr3R) was consistently enriched for all the different types of variants in offspring CTR samples (Supplemental Fig. S5). The genomes of offspring CTR flies contain the third chromosome balancer TM6B (*In(3LR)TM6B*), carrying multiple inversions to avoid recombination. These results suggest that the presence of the balancer induces an increased detection of variants by software based on the identification of split and discordant reads on this chromosome. Therefore, we recalculated the normalized number of SVs, SNP-indels and TE ISs considering exclusively the variants on chromosomes 2L, 2R, 4 and X. Results showed significant difference for BND at 10-12d stage and for SNP-indels at all stages (Fig. 5). Intriguingly, the number of SNP-indels resulted significantly higher in 0-2d HD than 0-2d CTR sample while at larval and 10-12d stages the trend was the opposite with a higher number of SNP-indels in CTR than HD brains. This preliminary observation has to be furtherly investigated and validated. Nevertheless, given that SNP-indels are enriched in 0-2d but decreased in 10-12d HD flies it might be tempting to hypothesize that they accumulate in cells that undergo cell death and are absent in aged, diseased individuals.

**Figure 5.**
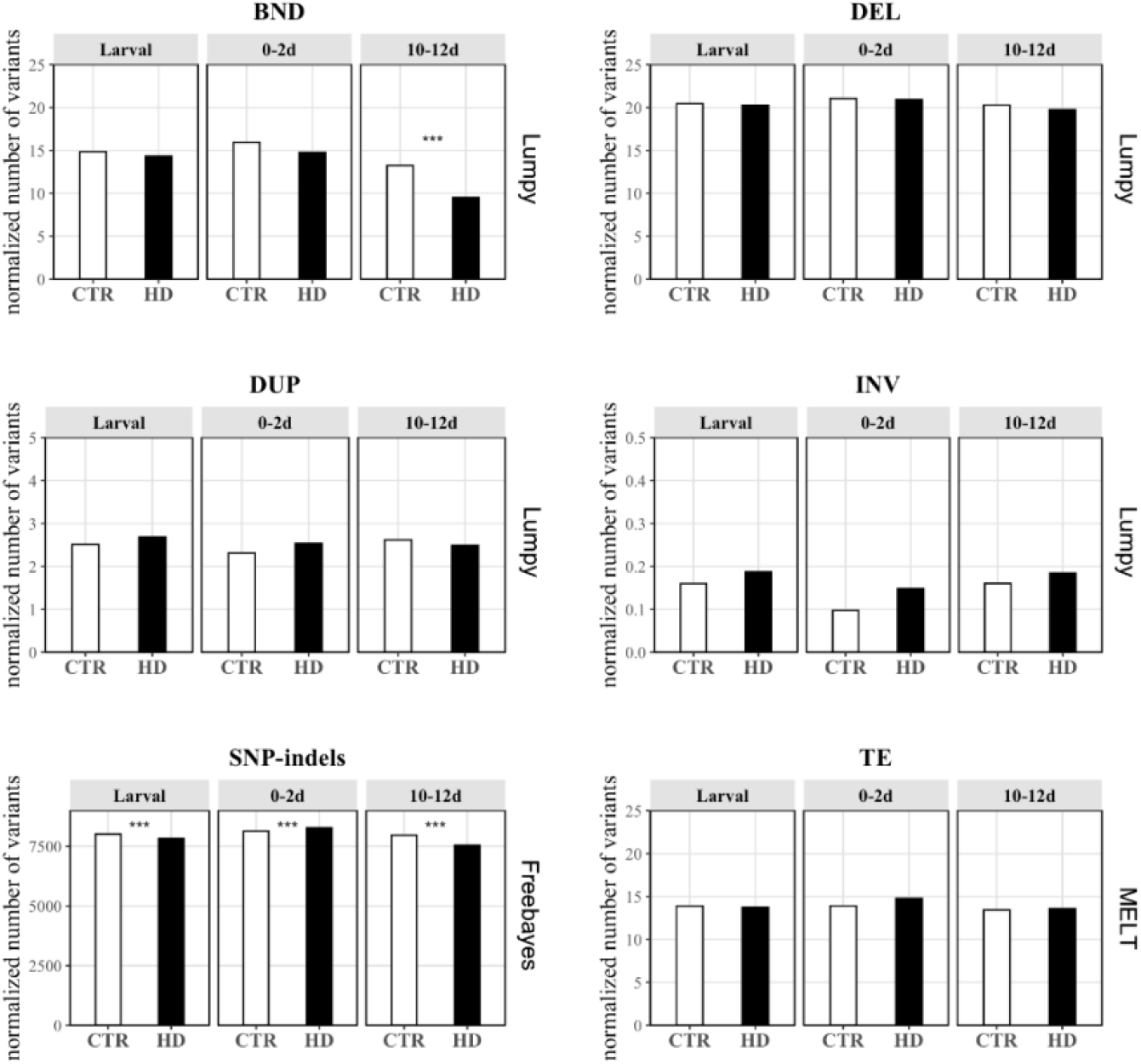
Number of analysed structural variants on chromosomes 2L, 2R, 4 and X. Break ends (BND), deletions (DEL), duplications (DUP), inversions (INV), SNP-indels and TE non reference ISs have been detected in each of the 6 samples (CTR and HD in larval, 0-2d and 10-12d time points) analysed in all chromosomes but no 3L and 3R. No changes in variants content were observed except for BND at 10-12d stage and SNP-indels showing an increased number in HD than CTR at 0-2d and a decrease in HD than CTR at both larval and 10-12d stages. The raw number of variants has been normalized on the total number of sequenced reads and multiplied by 1,000,000. Control samples are shown in white and HD in black. (FDR < 0.05, ** FDR< 0.01, *** FDR< 0.001, Two proportions Z-test with Benjamin-Hochberg FDR correction).

### Quantification of TE content by analysis of WGS data and validation by TaqMan CNV assays

The comparison of *de-novo* insertion sites by MELT did not show any significant difference. This analysis might be considered rather stringent since MELT, for technical reasons, discards TE ISs identified within a reference element (*i.e.* nested insertions) that account for an important fraction of novel insertions, especially in neurodevelopmental disorders (Jacob-Hirsch et al. 2018). We therefore developed an alternative bioinformatic strategy to quantify the overall TE content of our samples avoiding calling ISs on the exclusive basis of split and discordant read pairs. DNA-seq reads mapped on each TE consensus sequence were thus counted and the normalized values of these coverages were interpreted as the sample content for each TE. By analysing the coverage proportions of 189 TE classes, we highlighted 100 TEs showing a read coverage significantly different between CTR and HD samples at the pathogenic stage (0-2d stage) (FDR < 0.001) (Supplemental Table S1). Almost two third of them (65%) showed a higher content in HD samples. Fig. 6A presents data on the top 15 elements according to their statistical significance at 0-2d stage. Among them, 7 LTR retrotransposons (DM176, BEL_I, HMSBEAGLE_I, Gypsy2_I, STALKER-3, GYPSY_I, and BURDOCK_I) and 3 LINE retrotransposons (R1, I and DMRT1B) displayed a higher proportion of mapped reads in HD than CTR sample while the DNA element HOBO, 3 LTRs (GYPSY12_I, COPIA_I and ROO_I) and the LINE HET-A showed an opposite trend. The I-element is the LINE element displaying the highest ratio between mapped reads in HD and CTR samples at the pathogenetic stage (0-2d). GYPSY_I displayed the highest number of mapped reads in HD than CTR samples, in all the 3 analysed time points.

**Figure 6.**
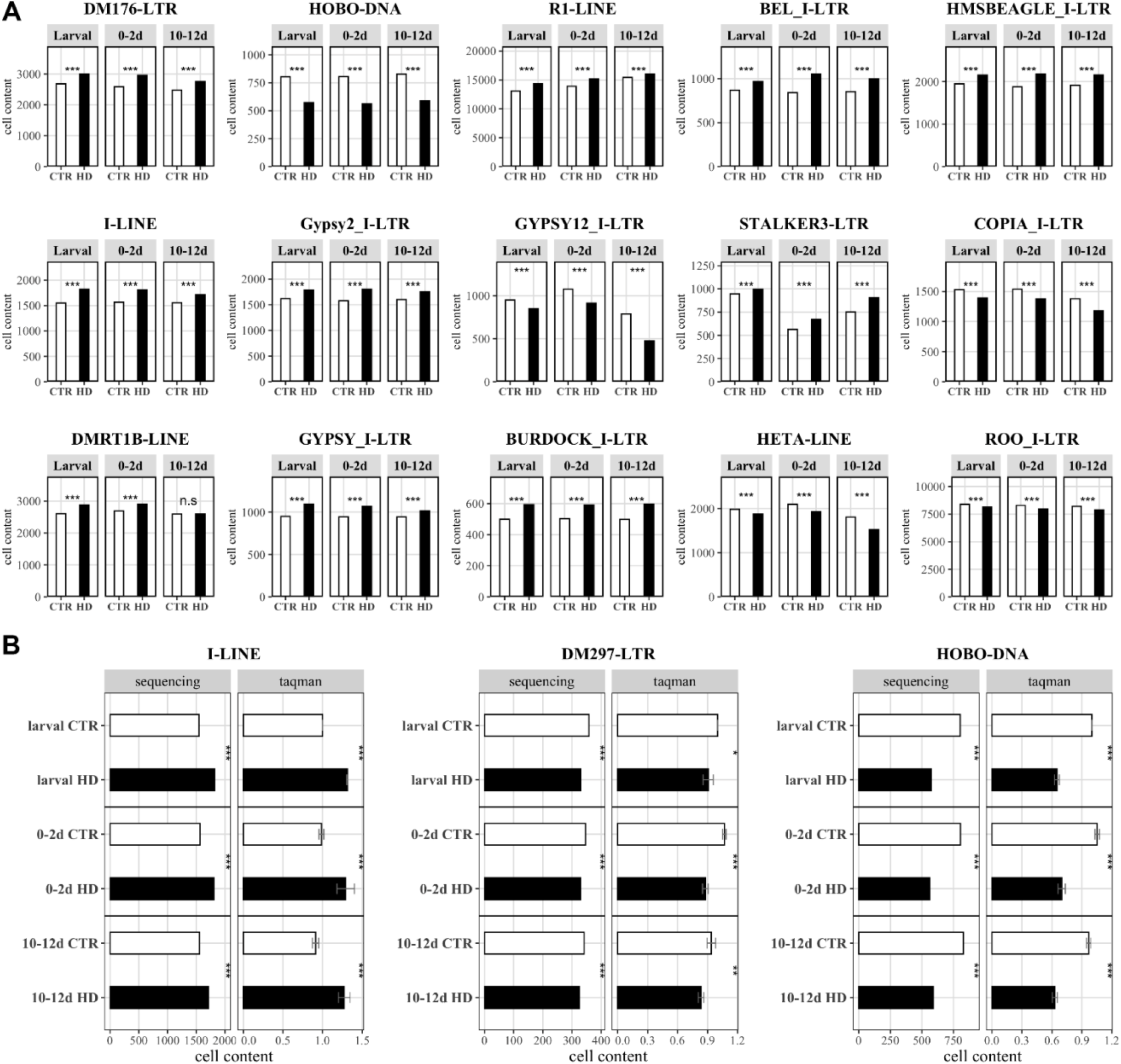
TE copy number according to sequencing analyses and TaqMan CNV assay. (A) 15 top significant TEs at the 0-2d stage. Cell content of each TEs has been calculated at larval, 0-2d and 10-12d for both CTR and HD samples. The top15 significant TEs at 0-2d stage (the pathogenetic one) are here represented. All the top15 TEs are classified as retrotransposons (LINE and LTR elements) except for HOBO, a DNA transposon. Of the 15 reported TEs, ten showed a higher cell TE content in HD than CTR 0-2d sample (DM176, R1, BEL_I, HMSBEAGLE_I, I-element, Gypsy2_I, STALKER-3, DMRT1B, GYPSY_I and BURDOCK_I). The cell TE content reported on y-axis has been calculated counting the reads mapping against each TE RepBase consensus and normalizing on the total number of sequenced reads and multiplied by 1,000,000. (*** FDR< 0.001, two proportions Z-test with Benjamin-Hochberg FDR correction). (B) Validation of the TE copy number by TaqMan CNV assay. The I-element (retrotransposon) assay fully confirmed the sequencing results highlighting higher content of I-element in HD samples than CTR samples, in all the 3 time points (left). TaqMan CNV assay and sequencing analyses showed concordant significant results for DM297 element (middle) (retrotransposon). TaqMan CNV assay displayed a lower CNV of HOBO (DNA transposon) in HD than control samples in all the 3 time points confirming sequencing results (right). Sequencing panel: cell content indicated as number of reads mapped on the TE consensus normalized on the total number of sequenced reads multiplied by 1,000,000. Two proportions Z-test with Benjamin-Hochberg FDR correction. Taqman panel: cell content reported as ddCt using as reference gene DMRT1C and as calibrator offspring larval CTR sample. (* p < 0.05, ** p < 0.01, *** p < 0.001, Two Way Anova with Bonferroni post hoc test). Offspring CTR samples are depicted in white whereas offspring HD in black.

To validate WGS data, we carried out a series of copy number variation (CNV) analysis by TaqMan quantitative PCR (qPCR) of the very same genomic DNA samples. While CNV Taqman assays are available for some TE classes in human and mouse genomes, a comprehensive toolbox is still lacking in *Drosophila*. Firstly, a properly designed TaqMan assay requires a normalizer. To this purpose we identified a specific, invariant DMRT1C fragment sequence that was stable in all samples and it presented a number of copies comparable with TEs (25 cycles threshold according to the 2-ΔΔCt method for relative quantification) (Livak and Schmittgen 2001). Then, we designed and optimized CNV Taqman assays for one representative class of each type of the following TEs: LINEs (I-element), LTRs (DM297) and DNA transposons (HOBO). As shown in Fig. 6B, TaqMan CNV assays confirmed the differential genomic TE content as measured by bioinformatic analysis of WGS data. When *actin* was used as invariant control in selective assays, results were comparable (data not shown).

### Preliminary evaluation of AZT treatment effects on TEs tissue content

We have previously shown that AZT treatment of HD flies rescued the eye phenotype and the genomic instability observed in larval brains and increased lifespan when administered at 0-2 days. Therefore, we investigated whether AZT affects the brain content of selected TE classes during the same time window when it exerts its systemic protective effects. As previously reported, we crossed parental lines on normal medium and transferred the HD offspring on AZT supplemented medium, starting the treatment at 0-2 or 4-5 days following eclosion. 10-12-day old flies were then harvested and brain tissue isolated. For each time point, (0-2 days – dark grey and 4-5 days – stripped bars), 3 adult brain gDNAs (HD untreated, HD treated with ethanol (solvent) and HD treated with AZT) were analysed and compared to gDNA from a healthy control with similar age (10-12 days old) (white). By taking advantage of Taqman CNV assays on arbitrarily selected TE classes, we monitored the tissues content of LINE I-element, LTR DM297 and DNA transposon Hobo. As shown in Supplemental Fig. S6, AZT treatment had no effects on tissue content of I-element (Supplemental Fig. S6A). This result suggests that the I-element activity might have started before AZT treatment at 0-2 days. Additionally, AZT treatment had no effects on the tissue content of the Hobo element (Supplemental Fig. S6C). This result was expected, as Hobo is a DNA transposon whose transposition mechanism does not exploit the reverse transcriptase enzyme thus not being affected by the AZT treatment. On the contrary, AZT decreases DM297 in a statistically significant manner when administered at 0-2d suggesting that AZT treatment may have impaired DM297 mobilization in this time window (Supplemental Fig. S6B).

### Identification of ISs

To investigate whether functionality of specific genes may be altered by TE ISs, we took advantage of TE IS coordinates predicted by MELT. Considering all the 6 samples, MELT identified 9,023, non-annotated ISs with ‘PASS’ quality score and supported by at least 1 read at left and 1 at right breakpoints. The 9,023 total nonannotated ISs correspond to 2,404 ISs non-redundantly represented among the 6 samples (ISs identified among different samples have been clustered together when closer than 50 nt). The majority of the non-reference TE ISs resulted to hit intronic and intergenic regions (51% and 39% of the total number of non-redundant ISs, respectively) with a small fraction of them affecting 3’ and 5’ UTRs and coding regions (6%, 2% and 2%, respectively). Gene ontology (GO) enrichment analysis performed on the 959 non redundant genes associated to the 2,404 TE ISs highlighted positive enrichments of GO terms mainly associated to development and neuronal functions such as synapses formation and signal transduction (Supplemental Table S2). Of the 189 TE class analysed, 129 of them present at least one event. Supplemental Table S3 presents the number of ISs for each TE class for all samples.

## Discussion

Growing evidence suggests a detrimental effect of TEs activity in neurodegenerative disorders (Tam et al. 2019). Taking advantage of a fly model, this study proposes a pivotal role played by TEs in HD pathogenesis. We provided in vivo functional evidence that pan-neuronal expression of 128Q mutant Htt in *Drosophila* induces a widespread deregulation of TE silencing and the activation of their expression. We found that different families of TE transcripts were significantly upregulated in larval, young (0-2 days old) and aged (10-12 days old) HD brains. Moreover, the increase of TE transcripts at 10-12 days correlated with age-dependent HD neurodegeneration. Interestingly, we found that TE upregulation in response to 128Q mutant Htt expression is not a specific feature of postmitotic neurons. Indeed, the expression of the pathogenic Htt protein in glial cell lineage by using the pan-glial driver repo-Gal4, induced similar altered levels of TE transcripts.

On the basis of previous studies documenting a global constitutive heterochromatin relaxation in neurodegenerative disorders (Frost et al. 2014) and identifying constitutive heterochromatin as an epigenetic silencing mechanism for TE (Sun et al. 2018; Fedoroff 2012), we investigated the underlying molecular mechanisms of HD-induced TE activation and demonstrated that the loss of TE silencing in 10-12 days old HD fly brains is supported by a significant reduction of H3K9me3 occupancy in their promoters and coding regions. Moreover, PEV assay on larval brains expressing HD pathogenic construct, confirmed that the overall reduction in H3K9me3 levels on TE sequences results from a global heterochromatin relaxation that impairs its ability to silence the variegating reporter transgene Hsp70-lacZ into Y pericentromeric heterochromatin (Lu et al. 1996). This is consistent with the observation that HP1 overexpression partially rescues the severe neurodegenerative phenotype induced by eye-specific expressions of the 128Q mutant Htt. In order to discriminate if the release of transposon silencing in HD is a cause or a consequence of neurodegeneration, we adopted a pharmacological approach to block the transposition of RTEs with RT inhibitors and demonstrated that RT inhibitors treatments rescued i) the HD eye phenotype, ii) the genomic instability observed in HD larval brains and iii) the lifespan in HD flies treated from larvae and 0-2 days adult stage thus suggesting that the HD phenotype may depend on RT activity and that retrotransposition at 0-2d stage disease actively contributes to neurotoxicity. Additionally, given the phenotypic rescues highlighted by the RT inhibitor treatments, it is likely that DNA transposons, whose transposition is not mediated by RT, do not contribute to neurotoxicity.

In the second part of our study we explored the genomic complexity of TE mobilization by WGS and Taqman qPCR. To this purpose we sequenced gDNA from HD brain tissues (*elav-Gal4>128QHtt*, offspr. HD) and offspring control (*elav-Gal4/+*, offspr. CTR) at 3 different developmental time points (larvae, young and aged adults). We then carried out an extensive bioinformatic analysis of their differences in TE content respect to the genome of reference. This approach presents the limitations associated to sequencing “bulk” DNA from 60 individuals and from samples composed by a highly heterogeneous pool of different cell types. Analysis of cellular mosaicism in mammalian brains have led to a wide range of estimates of novel insertions per cell (from 0.2 to 16.3) (Evrony et al. 2016; Upton et al. 2015). The representation of these relatively rare somatic ISs in WGS data strictly depends on the time of the retrotransposition event during embryo development or during the lifespan of an individual and the consequent number of genomes that contain a single integration. Given the depth of sequencing and the heterogeneity of the sample we are conscious that this analysis underestimates the number of somatic ISs unveiling only germinal or mobilization events that took place very early in embryogenesis. Despite these limitations we were able to identify a total of 9,023 non-reference ISs with ‘PASS’ quality score. GO enrichment analysis showed that ISs were mainly associated to development and neuronal functions such as synapses formation and signal transduction, well-known targets in HD pathogenesis.

When measuring the number of SVs for each class of them, we confronted another confounding factor due to the presence, in the CTR genomes, of the third chromosome balancer TM6B (*In(3LR)TM6B*) carrying multiple inversions to avoid recombination. The presence of this highly rearranged chromosome resulted in an increased detection of SVs by softwares based on the identification of mismatches for SNPs call and of split and discordant reads for non-reference TE ISs call. To normalize for this effect, we recalculated variants leaving out data from this chromosome. Nevertheless, we found that a specific type of variants, SNP-indel, was differentially present in all HD genomes. Given that SNP-indels are enriched in 0-2d but decreased in 10-12d HD flies we may speculate that they accumulate in cells that undergo cell death and are absent in aged, diseased individuals.

We then developed a complementary bioinformatic approach based on assessing the normalized number of reads that map on the consensus sequence for each of the 189 TE classes in the *Drosophila melanogaster* genome. This approach avoids relying exclusively on split and discordant reads. We thus identified 100 TE class which sequence content was different between HD and CTR flies at 0-2d, the time when systemic RT inhibitory treatment was increasing lifespan. Importantly, this differential genomic content was confirmed with Taqman qPCR. While data from sequencing coverage and qPCRs may suffer from differential access to selective regions of the genome, a very good correspondence was evidenced between the two experimental approaches. Moreover, this analysis identified a list of TE classes with a differential tissue content and therefore candidates for being molecular effectors of RT-dependent phenotypes. Among them, LTR DM297 copy number decreased in the presence of AZT at 0-2d of HD brains enlisting this TE class for further analysis.

A minority but substantial fraction of TEs showed a decrease in HD samples. This is in agreement with recent evidence that TE mobilization may also lead to genomic deletions (Rodriguez-Martin et al. 2020).

As previously reported in other biological systems, a lack of correlation between expression and mobilization was evident and probably due to the role of TE RNAs in chromatin remodelling independent from retrotransposition (Wang et al. 2018; Jachowicz et al. 2017).

Interestingly, GYPSY_I showed both a strong increase of RNA expression levels and the highest number of mapped reads in HD than CTR samples, in all the 3 analysed time points. GYPSY_I element was previously described (Krug et al. 2017) to be over-expressed and contribute to neurodegeneration in a *Drosophila* ALS model (Krug et al. 2017). Furthermore, Gypsy expression was found induced in a fly model of tauopathy (Guo et al. 2018). Our results suggest that GYPSY_I may be involved in three different fly models of neurodegenerative diseases such as ALS, AD and HD.

In summary, the genomic analysis of TE content in HD fly brains lays down the foundations for the identification of the TE classes causally involved in the establishment of RT-dependent HD phenotypes and for new strategies of pharmacological intervention.

## Methods

### *Drosophila* strains

The *Drosophila* stocks used in this study were obtained from Bloomington Drosophila Stock Center (Indiana University, Bloomington, IN) or Vienna Drosophila Resource Center (VDRC, www.vdrc.at) and are listed below:

*w^1118^; P{UAS-HTT.128Q.FL}f27b* (#33808); *w^1118^; P{UAS-HTT.16Q.FL}F24/CyO* (#33810)

*y^1^, w*; Dp(3;Y)BL2, P{HS-lacZ.scs}65E* (#57371); *w^1118^; P{GD12524}v31994*.

The Gal4 lines used in this study were: elav-Gal4 (*P{GawB}elav^C155^*, #458); repo-Gal4 (*w^1118^; P{GAL4} repo/TM3*, *Sb*^1^, #7415); GMR-Gal4 (*w*; P{GAL4-ninaE.GMR}12*, #1104).

The UAS-HP1a/CyO strain overexpressing HP1a under the control of UAS sequence was generated in our lab.

The Ore-R stock and balancer stocks, used to balance inserts on the X, second, and third chromosomes, respectively, have been kept in our laboratory for many years.

All flies were raised at 24 °C on standard cornmeal-sucrose-yeast-agar medium.

All crosses were performed at 24°C unless otherwise noted.

### NRTI food preparation

Zidovudine (AZT; PHR-1292 Sigma-Aldrich) and Lamivudine (3TC; PHR-1365 Sigma-Aldrich) were dissolved in 95% ethanol and water respectively. NRTI solutions were added in standard medium (cornmeal-yeast-sucrose-agar) to give a final concentration of 5 mg/mL for AZT and 1 mg/mL for 3TC.

### Mitotic chromosome preparations

Cytological preparations of mitotic chromosomes from *Drosophila* larval brain were obtained according to Pimpinelli et al. (2000) (Pimpinelli et al. 2000) and stained with DAPI (4,6-diamidino-2-phenilindole, 0.01 mg/ml) to visualize DNA.

The slides were mounted in antifading medium (23,3 mg/mL of DABCO (1,4-Diazobicyclo-(2,2,2) octane) in 90% glycerol – 10% 1X PBS) and all images were acquired on Ellipse Epifluorescence microscope (E1000 Nikon) equipped with a CCD camera (Coolsnap). Images were analysed and further processed using Adobe Photoshop CS6.

### Immunofluorescent staining of *Drosophila* larval brains

The immunofluorescent staining of larval brains was performed according to Wu & Luo (2006) (Wu and Luo 2006). The primary antibody used was rabbit anti-γH2AV (1:50, 600-401-914 Rockland). Fluorescent labelled secondary antibody raised in goat was Alexa Fluor 555 conjugated anti-rabbit (1:300, A21437 Thermo Fisher Scientific). The brains were stained with TOTO-3 Iodide (1 μM) to visualize DNA and mounted in antifading medium. Confocal observations were performed using a Leica DMIRE (Leica Microsystems, Hiedelberg, Germany) and a Zeiss LSM 780 (Zeiss, Berlin, Germany). Images were analysed and further processed using Zen Software and Adobe Photoshop CS6.

### *Drosophila* eye imaging

Age- and genotype-specific flies were ether-anesthetized and eye external images were taken with a Nikon camera D5000 mounted onto a stereoscopic microscope. Images were further processed using Adobe Photoshop CS6. The *Drosophila* eye defects were quantified using Flynotyper (Iyer et al. 2016), a software that detects alterations in eye morphology and that provides a Phenotypic Score as a quantitative measure of the severity of the ommatidial arrangement defects. Data was then analysed with ANOVA and Tukey’s multiple comparison tests.

### Lifespan Analysis

150-200 flies were collected for each experimental group (30 flies per vial). Every 2–3 days, flies were passed into new vials and dead flies were counted. The survival rate was calculated by the percentage of total flies surviving.

### Total RNA extraction and qRT-PCR

RNA samples from larval and adult brains were isolated using Qiazol reagent (Qiagen), according to the manufacturer’s instructions. 5 μg of total RNA was reverse transcribed using oligo dT and SuperScript Reverse Transcriptase III (Invitrogen) according to the manufacturer’s protocol. The qPCR reactions were carried out with QuantiFast SYBR Green PCR Kit (Qiagen) according to manufacturer’s protocol. Relative abundance of the different transcripts was determined using the 2^-ΔΔCt^ method (Livak and Schmittgen 2001) using rp49 or gapdh transcript as controls. qRT-PCR experiments were performed in three independent biological replicates each with three technical replicates. All primer used are listed below:

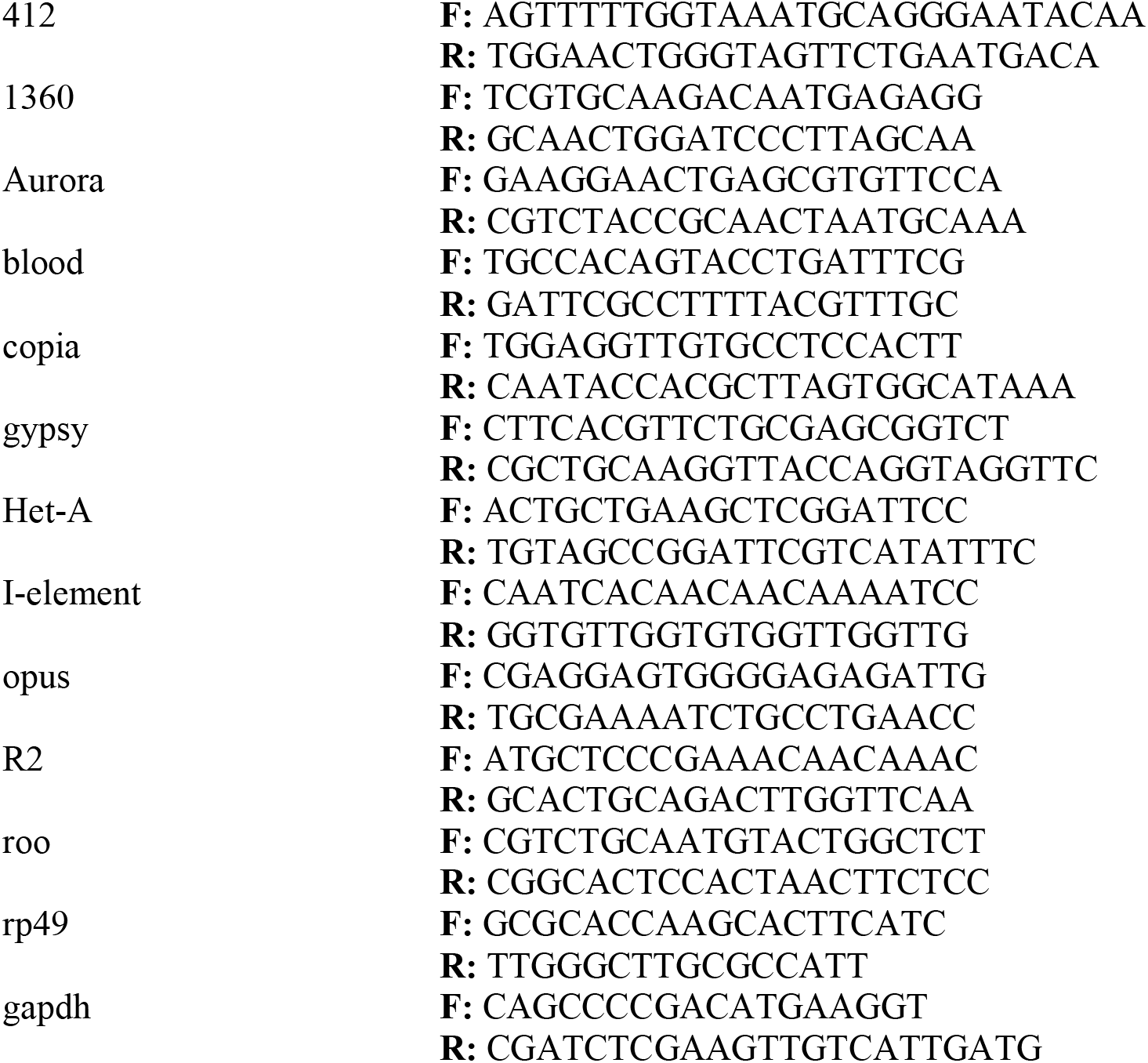

### Semi-quantitative PCR

About 100ng of reverse transcribed total RNA purified from fly heads, was added to the following reaction mix: 1X PCR Buffer, primers (0.2 μM), MgCl_2_ (1.5 mM), dNTPs (0.2 mM each), and 2U/rxn of Platinum™ Taq DNA Polymerase (Invitrogen). The thermal profile was: 94 °C for 3 minutes for the initial denaturation, followed by 30 cycles at 94 °C for 30 seconds, 60 °C for 30 seconds, 72 °C for 30 seconds and a final extension at 72 °C for 7 minutes. PCR products were analysed by agarose gel electrophoresis.

The oligonucleotides used as primers were:

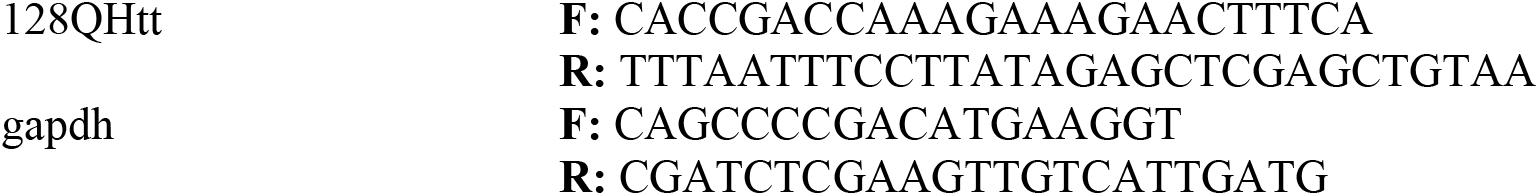

### Western blot analyses

Western blot was carried out as previously described in Cappucci et al., (2019) (Cappucci et al. 2019). To obtain a total protein extract, *Drosophila* brain or head samples were homogenized in Sample Buffer 1X (50 mM Tris-HCl pH 6.8, 100 mM dithiothreitol, 2% sodium dodecyl sulphate, 2.5% glycerol, 0.1% bromophenol blue) and heated at 85 °C for 8 minutes to complete the protein denaturation.

Protein extracts were fractionated by 10% SDS-PAGE and electroblotted onto Immobilion-P polyvinyl difluoride membranes (Bio-rad) in CAPS-based transfer buffer (10 mM CAPS pH 11, 10% methanol) in a semi-dry transfer apparatus (Amersham Biosciences). The membranes were blocked with 5% non-fat dry milk in Tris-buffered saline with Tween 20 (TBST) buffer (20 mM Tris pH 7.5, 150 mM NaCl, 0.1% Tween 20) and incubated with the following antibodies diluted in TBST: mouse anti-ENV 8E7 (1:500, kindly provided by J. Gall), rabbit anti-GIOTTO (1:10000, kindly provided by G. Cestra), mouse anti-elav (1:500, 9F8A9 DSHB) and mouse anti-repo (1:500, kindly provided by G. Cestra). Proteins of interest were detected with HRP-conjugated goat anti-mouse or anti-rabbit IgG antibody and visualized with the ECL Western blotting substrate (GE Healthcare), according to the provided protocol. The chemiluminescence detection was performed on the ChemiDoc XRS+ System (Biorad) and analysed using the included ImageLab software.

### Chromatin immunoprecipitation assay (ChIP)

Chromatin immunoprecipitation was performed according to Casale et al., (2019) (Casale et al. 2019) with some modifications. Approximately 250 μL of 10-to 12-days-old fly heads were homogenized in 3 mL of NEB buffer (10 mM HEPES-Na pH 8, 10 mM NaCl, 0.1 mM EGTA Na pH 8, 0.5 mM EDTA-Na pH 8, 1 mM DTT, 0.5% NP-40, 0.5 mM Spermidine, 0.15 mM Spermine, 1× EDTA-free Complete Protease Inhibitors) with a Polytron homogenizer (Kinematica Swizerland) with a PT300 tip five times (2 min and 1 min on ice) at 3500 rpm and five times at 4000 rpm. The homogenate was transferred to a pre-chilled glass dounce (Wheaton) and 20 full strokes were applied with a tight pestle. Free nuclei were filtered on a 70 μM strainer and then centrifuged at 6000xg for 10 min at 4 °C. The nuclei-containing pellets were resuspended in 1 mL of NEB + 0,25 M sucrose and centrifuged at 20000xg for 20 min on sucrose gradient (0.65 mL of 1.6 M sucrose in NEB, 0.35 mL of 0.8 M sucrose in NEB). The pellet was resuspended in 1 mL of NEB and formaldehyde to a final concentration of 1%. Nuclei were crosslinked for 10 min at room temperature and quenched by adding 1/10 vol of 1.375 M glycine. The nuclei were collected by centrifugation at 6000xg for 5 min. Nuclei were washed in PBS 1X and then twice in 1 mL of NEB and resuspended in 1.5 mL of Lysis Buffer (15 mM HEPES-Na pH 7.6, 140 mM NaCl, 0.5 mM EGTA, 1 mM EDTA pH 8, 1% Triton X-100, 0.5 mM DTT, 0.1% Na-Deoxycholate, 0.1% SDS, 0.5% N-lauroylsarcosine and 1× EDTA-free Complete Protease Inhibitors). Nuclei were sonicated using a Hielscher Ultrasonic Processor UP100H (100W, 30kHz) thirty times for 30s and 30s on ice. Sonicated nuclei were centrifuged at 15000xg for 10 min at 4 °C. The majority of sonicated chromatin was 300 to 500 base pairs (bp) in length. To allow the immunoprecipitation, 3μg of H3K9me3 monoclonal antibody (61013 Active Motif) were incubated in the presence of dynabeads protein G (Invitrogen) for 3 h at room temperature on a rotating wheel, then chromatin extract was added and incubation was continued overnight at 4 °C on a rotating wheel. The supernatants were discarded and samples were washed twice in Lysis Buffer (each wash 15 min at 4 °C) and twice in TE Buffer (1 mM EDTA, 10 mM Tris HCl pH 8). Chromatin was eluted from beads in two steps; first in 100 μl of Elution Buffer 1 (10 mM EDTA, 1% SDS, 50mM Tris HCl pH 8) at 65 °C for 15 min, followed by centrifugation and recovery of the supernatant. Beads material was re-extracted in 100 μl of TE + 0.67% SDS. The combined eluate (200 μl) was incubated overnight at 65 °C to revert the cross-link and treated by 50 μg/ml RNaseA for 30 min at 37 °C and by 500 μg/ml Proteinase K (Invitrogen) for 3 h at 65 °C. Samples were phenol-chloroform extracted and ethanol precipitated. DNA was resuspended in 15 μl of water and candidate genes were amplified through qPCR. The DNA oligonucleotide primers needed for ChIP-qPCR are listed below:

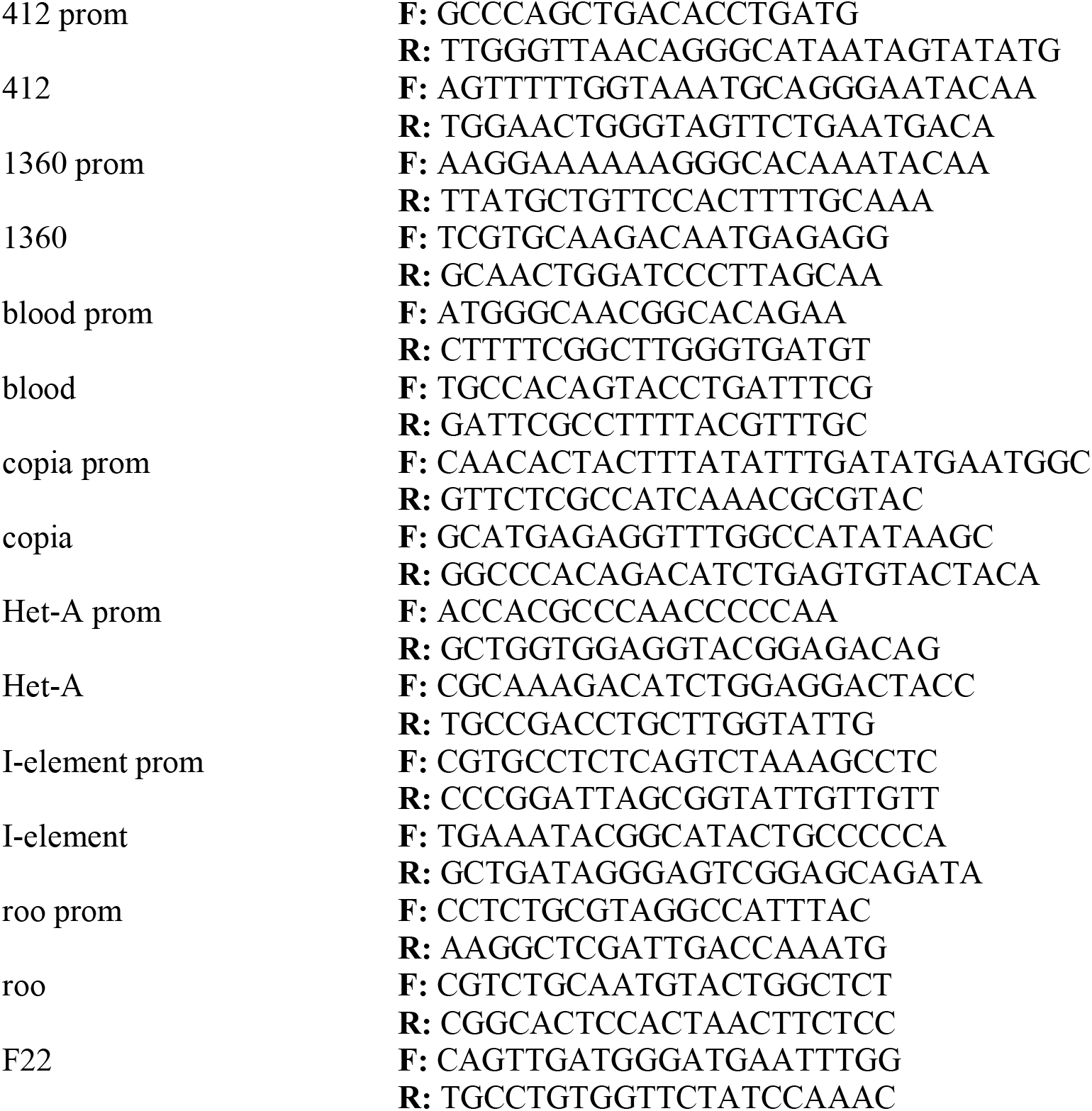

### Histochemical localization of β-galactosidase in PEV analysis

The induction and localization of β-galactosidase on larval brains was performed according to Lu et al., (1996) (Lu et al. 1996). To induce β-galactosidase expression, larvae were heat-shocked at 37 °C for 45 minutes followed by 1 hour of recovery at room temperature. Brains were dissected in 1X PBS, fixed in 4% formaldehyde for 20 minutes, washed in 1X PBS and incubated in 0.2% X-gal (5-bromo-4-chloro-3-indolyl-p-D-galactopyranoside) assay buffer (3.1 mM potassium ferricyanide, 3.1 mM potassium ferrocyanide, 10 mM PB (pH 7.2), 0.15 M NaCl, 1 mM MgCl2) for 16 hours. Brains were immersed in 1X PBS and the images were acquired through a stereomicroscope equipped with a Nikon D5000 camera and further processed using Adobe Photoshop CS6. X-Gal staining intensities were analysed using ImageJ software. All RGB images were converted in 8-bit grayscale and then inverted with the command *Edit>Invert*. An outline was drawn around each brain, area and intensities measured, along with adjacent background reading. Intensity values were then calculated following the total corrected cell fluorescence (TCCF) method as described in McCloy et. al (2014) (McCloy et al. 2014). Statistical analysis (unpaired Student t test) was performed using GraphPad Prism 6 software.

### Isolation of genomic DNA

Brain tissues were homogenized in 200 μl of extraction buffer (120 mM Tris-HCl pH 8, 60 mM EDTA pH 8, 80 mM NaCl, 160 mM sucrose, 0.5% v/v SDS, 200 μg/mL RNase DNase free) and incubated at 65°C for 60 min. After cooling at room temperature for a few minutes, 28 μl 8M K-acetate was added. After 30 min on ice, the samples were spun at 10000 rpm for 15 min at 4°C; DNA was precipitated by adding 0.5 volume of isopropanol to the supernatant, leaving for 10 min at room temperature, and spinning again for 10 min. The pellet was washed with 70% ethanol, dried and redissolved in 60 μl H_2_O. 1 μg of gDNA was sequenced by high-throughput Illumina technology.

### DNA sequencing and raw reads quality controls

1μg of DNA was extracted from every fly sample. Sample1, sample3 and sample5 were obtained from homogenized brain tissue from offspring control flies (*elavG4/+; +/TM6*) at larval, young (0-2 days old) and aged (10-12 days old) stages while sample2, sample4 and sample6 were obtained from offspring HD flies (*elavG4/+; 128QHtt/+*) at the 3 different time points used for control samples. DNA was sequenced by Institute of Applied Genomics (IGA, Udine, Italy) obtaining 125 bp paired-end (PE) reads. The quality of the raw reads was tested using FastQC (version v0.11.7).

### Genomic structural variants analysis

For each of the 6 samples raw reads were aligned to the fly reference genome (dm6 release from UCSC database) with bwa (Li and Durbin 2009) using standard parameters, duplicated reads were marked and split-reads and discordant reads were extracted with samblaster (Faust and Hall 2014). Finally, the alignment files were sorted and indexed with Sambamba (Tarasov et al. 2015). The bam files obtained from the alignment were used for variant discovery, which we performed with the softwares Freebayes (Garrison and Marth 2012) and Lumpy (Layer et al. 2014).

Freebayes is a genetic variant detector designed to find small polymorphisms, such as SNPs (single-nucleotide polymorphisms), indels (insertions and deletions), MNPs (multi-nucleotide polymorphisms), and complex events (composite insertion and substitution events) smaller than the length of a short-read sequencing alignment. Freebayes was run on the aligned reads with standard parameters (minimum variant QUAL score to output = 1); and produced a vcf file version 4.2 as output. Only variants featuring a QUAL score above 20 were considered as putative variants. Lumpy is a structural variant (SV) discovery framework that integrates data from depth coverage, split-reads and discordant pairs from an aligned genome. It is able to detect inversions (INV), duplications (DUP), deletions (DEL), and break ends (BND). Structural variants genotyping has been performed with the tool SVTyper (Chiang et al. 2015). Lumpy was run on the aligned reads with standard parameters (minimum sample weight for a call = 4, trim threshold = 0) and produced a vcf file version 4.2 as output. Only variants featuring a QUAL score above 20 were considered as putative structural variants. The raw number of structural variants, for every typology of variants, was normalized on the total number of sequenced reads and multiplied by 1,000,000.

### TE insertion sites analysis

In order to detect non-reference insertion sites (ISs) we run the Mobile Element Locator Tool (MELT) (version 2.1.4) (Gardner et al. 2017) in *Single* mode with the following parameters: *-c:* (coverage value) average coverage calculated with the pileup.sh script of the bbmap package (Bushnell B. - BBMap), *-h*: fly genome used for the mapping analysis (dm6 downloaded from UCSC database), *-bamfile*: the sorted bam alignment file of each sample, *-n*: bed file containing gene annotations for the dm6 fly genome retrieved from UCSC, *-t*: mobile element insertions (mei). To create the mei files, the MELT command *BuildTransposonZIP* was used providing, for each TE (having removed SAT and Unknown TES), the fasta sequence (downloaded from RepBase (Bao et al. 2015), the bed file containing the annotated genomic copies to mask during the insertional analysis and the error rate value that was set to 5 (maximum number of mismatches allowed per 100 bases of the MEI reference during alignment). After having run MELT on each of the 6 samples for all the fly TE consensus, the vcf output file was filtered in order to select ISs with the ‘PASS’ quality score and with at least 1 read supporting both left and right breakpoints. Then the number of ISs was normalized on the total number of sequenced reads and multiplied by 1,000,000.

### Gene ontology (GO) enrichment analysis

GO enrichment analysis has been performed by using topGO (Alexa and Rahnenfuhrer 2019) on GO terms associated to the genes harbouring at least one TE non-annotated IS and using as background list GO terms associated to the whole set of protein coding and non-coding annotated genes. To test for positive enrichments in genes harbouring TE non-annotated ISs with respect to background Fisher’s Exact Test (algorithm=‘weight’) has been used. GO terms associated to less than 10 significant genes have been discarded prior to FDR (Benjamini & Hochberg correction). Significant threshold has been imposed to FDR < 0.01.

### *In silico* TE tissue content quantification

In order to *in silico* quantify TE cell content in the 6 sequenced samples, DNA-seq reads of every sample were mapped on each of the Repbase (Bao et al. 2015) *Drosophila Melanogaster* TE consensus sequences (with exception for SAT and Unknown TEs) using bwa(Li and Durbin 2009) with default parameters (bwa version 0.7.15-r1140). Reads mapping on each TE consensus sequence were then counted using samtools view (-F 4 and -c parameters, version 1.3.1) (Li et al. 2009). Two proportions Z-test (prop.test function in R) was used at larval, 0-2d and 10-12d stages to highlight significant differences in the proportion of reads mapping on TEs in CTR and HD samples with respect to the total number of sequenced reads. P-value scores were then adjusted applying false discovery rate (FDR) correction (Benjamini & Hochberg) and the set of significant TEs was defined selecting TEs with FDR < 0.001 at 0-2d stage.

### DNA quantification for primers validation and CNV assays

In order to discriminate small variations in TE contents, a precise quantification of genomic DNA is required. The first DNA dilution to 2 ng/μL was spectrophotometrically measured using NanoDROP (Thermo Scientific, Thermo Fisher Inc, USA). Further dilutions in TE buffer (10mM Tris-HCl, 1 mM EDTA, pH 7.5) to a final concentration of 100 pg/μL were performed using QuantiT™ PicoGreenR dsDNA kit (Invitrogen, Thermo Fisher Inc, California, USA), following manufacturer’s instructions. Standard curve ranging from 20 pg/mL to 640 pg/mL, DNA samples and blank were measured in duplicate using EnSpire® Multimode Plate Reader (PerkinElmer Inc, Waltham, USA). By plotting the measured fluorescence versus DNA, concentrations were calculated through a standard curve plot. Final DNA concentrations ranging from 90 to 110 pg/μL were accepted for the PCR analysis.

### Primers and probes design and PCR amplification

Following standard experimental protocols for validation assay, primers and probes were designed on the candidate TE consensus sequences using the online tool Primer3 (https://primer3.org/) as listed below:

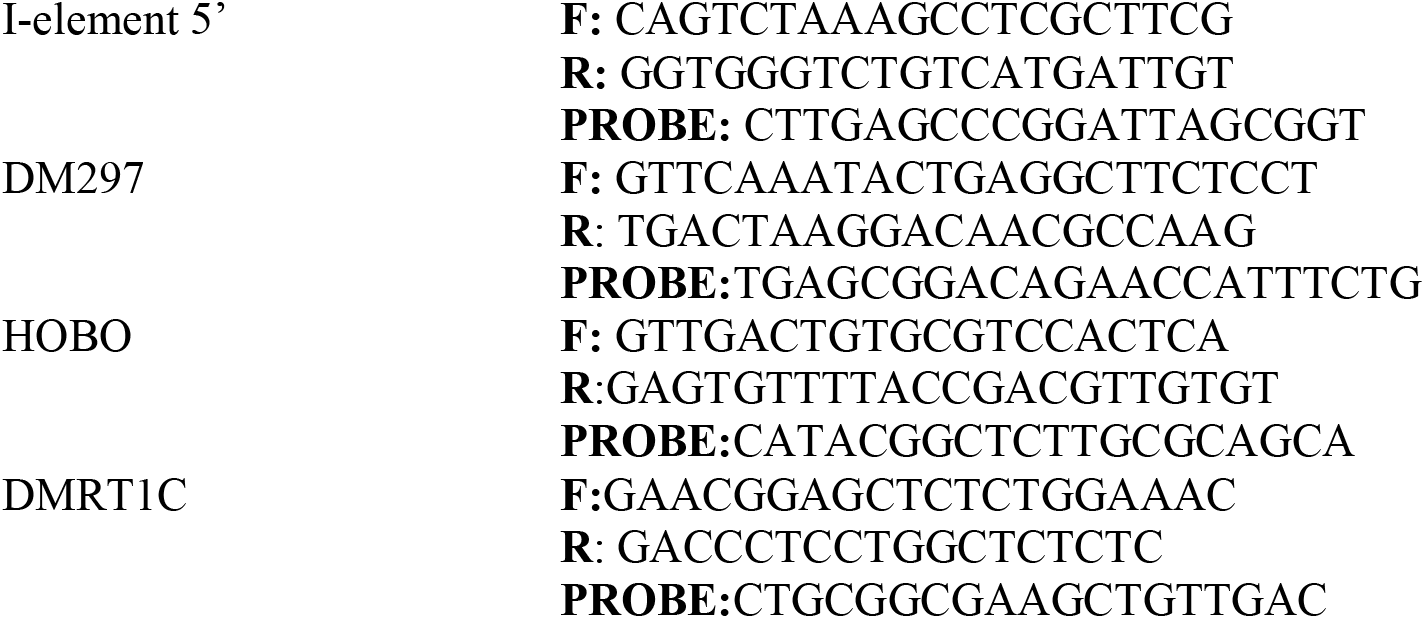

Primers and probes specificity was tested using an *in silico* amplification custom bioinformatics pipeline. Briefly, for each candidate TE, primers and probe were mapped using bowtie (version 1.2) (Langmead et al. 2009) against the dm6 fly genome allowing 0 mismatches in the last 7 nucleotides and up to 3 in the remaining part. Then only the *in silico* amplicons i) recognized by primers on opposite strand, ii) with the probe mapped in the between and iii) with a length comprised between 50 and 2000 nt, were selected. To test their specificity, the genomic coordinates of the selected *in silico* amplicons were intersected (intersectBed, -wo, -bed, -F 0.99 parameters (Quinlan and Hall 2010) with the coordinates of TEs annotated in the fly dm6 genome. The same custom pipeline was then re-run mapping primers and probes against the TE consensus sequences, in order to select primers and probe that recognize a unique TE sequence. Primers and probes were finally synthesized by Sigma-Aldrich. All primers were tested on 100 ng of genomic DNA in order to validate experimentally their specificity. PCR amplifications were performed using 5units/μL TaKaRa ExTaq Polymerase, 10x TaKaRa Buffer (both ClonTech), dNTPs (2,5 μM) and primers (10 μM each). PCR conditions are as follows: a denaturation step at 94°C for 3 minutes, followed by 40 cycles of amplification at 94°C for 30 sec, 60°C for 30 sec, 72°C for 1 minute, ending with a final extension at 72°C for 5 minutes. The amplified products together with a negative control, such as non-templates, were separated on a 1.5 % agarose gel and visualized with Midori staining. Amplicons were extracted from gel following manufacturer’s instructions (QIAquick Gel Extraction Kit, Qiagen) and confirmed through Sanger sequencing (Eurofins Genomics). For the DM297 LTR and Hobo DNA elements primer and probe have been designed against the 3’ portion of the consensus sequence while for I-element a design on the consensus 5’ has been performed.

### Invariant gene design for TaqMan CNV assay

In order to evaluate the number of copies of repetitive elements, a proper invariant gene should be a multi-copied genomic element. A TaqMan CNV assay requires an invariant gene resulting stable in all the samples analyzed and with a number of copies similar to the analyzed elements. To our knowledge, all the previous genomic studies performed with a TaqMan qPCR assay to estimate copy number variations of TEs in fruit flies, were performed using single-copy genes as invariant gene, such as *actin, gapdh, rp49 (rpl32)* (Krug et al. 2017; Guo et al. 2018). However, in our experimental conditions (100 pg DNA) and following Real Time qPCR technical protocols, single copy gene as actin amplified within 35 cycles threshold (Ct), resulting not comparable to the TEs (on average 26 Ct) in the final 2–ΔΔCt method for the relative quantification (Livak and Schmittgen 2001). To overcome this technical problem, we scanned the fly genome to find a stable multi-copied element. We thus took advantage of the IS analysis carried out with MELT on our WGS data considering that ancient, fixed and inactive TEs could represent candidate invariant sequences. 56 TE classes showed no novel ISs in all the 6 fly samples. Given that we are selecting TEs with no novel ISs, the balancer-associated TE ISs over-estimation in CTR samples should not affect this analysis. We then selected TEs with at least 30 annotated genomic copies and with a consensus sequence longer than 400 bp, shortening the list to 5 TEs. Among these, DMRT1C LINE element was the only element displaying no full-length copies fixed in the fly genome. The DMRT1C consensus sequence is, indeed, 5,443 bp long, whereas the length of the 224 DMRT1C fixed fragments in the fly genome is comprised between 30 and 3,000 bp. Additionally, none of the DMRT1C 224 copies retained a full-length reverse transcriptase functional domain. Together, these observations strongly suggest that all the 224 DMRT1C genomic copies are likely truly truncated and inactive. DMRT1C copy number *in silico* predicted, was further confirmed by PCR when compared to *actin* allowed the co-amplification with the other TEs under study at very low concentrations (100 pg), in optimal technical conditions (31 Ct). The DMRT1C was thus validated as the best candidate invariant element and used in TaqMan assays as normalizer (data not shown).

### Taqman based Copy Number Variation (CNV) assay

Taqman based Copy Number Variation (CNV) assay was performed to detect TEs’ content in DNA in drosophila samples in all timing and life conditions, as previously described. All custom TaqMan probes sequences are listed above and were designed following standard protocol and the vendor’s custom assay design service manual (Thermo Fisher Scientific). Probes for all TE assays were conjugated to the fluorophore label MGB 6-FAM, whereas the invariant genes *Actin* or DMRT1C probes were conjugated with MGB-VIC. Actin assay Act5C (assay ID Dm02361909_s1) was acquired from Thermo Fisher Scientific.

A 20 μl reaction volume containing 10 μl of 2X iQ Multiplex Powermix (Biorad), 0,60 μl of primers (10 μM), 0,2 μl of probes (10 μM) and 1 μl DNA (100 pg for DMRT1C; 1ng for Actin) was used for qPCR amplification. PCR conditions are as follows: 95°C for 20 sec, followed by 40 cycles of amplification at 95°C for 10 sec and 59°C for 30 sec.

Standard curves were performed for each couple of target and invariant gene on genomic DNA ranging from 32 pg to 2 ng. The slope of linear regression to standard curve was nearly −3.32 for all assays in this study, which means that primers and multiplexing efficiency were acceptable.

The experiments were carried out using CFX96 Real Time PCR detection system (Biorad), according to standard protocol. Three replicas of qPCR reactions were performed in duplicates. Standardization was performed considering a control sample as calibrator and an independent inter-run calibrator for each plate. Data obtained from the co-amplifications of the target DNA sequence and the internal invariable control were analyzed using the 2-ΔΔCt method for the relative quantification (Livak and Schmittgen 2001).

### Statistical analysis

Statistical analyses were performed using GraphPad Prism version 6.00 (GraphPad Software, La Jolla, CA, USA, www.graphpad.com). For all statistics a p value ≤ 0.05 was considered statistically significant. qRT-PCR, ChIP-qPCR and X-Gal Intensity statistical analysis was performed by the Unpaired t-test. Western blots were analysed by One-sample t-test. Lifespan data were analyzed by log rank test. Statistical significance of chromosomal abnormalities was determined by Chi-Square test. Mitotic index and Phenotypic Score (Flynotyper) statistical significance was determined by one-way ANOVA followed by Tukey’s post-hoc test. Statistical details and significance levels can be found in the figure legends. For the TaqMan CNV assay, to compare all study groups and experimental conditions, Two Way ANOVA with Bonferroni Post Hoc Test were used, as described specifically in each result. Statistical significance for all the results obtained from the bioinformatics analyses (number of variations with/without chr3 and TE cell content) was determined by using the two proportions Z-test (prop.test function in R). Calculated p values were then corrected using Benjamini & Hochberg FDR correction. FDR < 0.05 were considered as statistically significant.

## Data availability

The data that support the findings of this study are included in the article and/or the Supplementary Materials. Additional data related to this paper may be requested from the authors

## Acknowledgements

We are grateful to Sergio Pimpinelli for critically reading the manuscript. We thank the Bloomington Stock Center, VDRC Stock Center, and the Developmental Studies of Hybridoma Bank for fly strains and antibodies. We would like to thank Gianluca Cestra and Joseph Gall for providing giotto, repo and env antibodies. We sincerely appreciate Silvana Caristi for assistance with confocal image acquisition. We are indebted to Ms Eva Ferri, Fabrizio Torri and Alessandra Sanna (IIT), Cristina Leonesi (SISSA) for administrative support.

## Authors’ contributions

A.M.C., F.L., and U.C., performed the molecular and genetic experiments, contributed to the analysis of the results and to the design of the research; F.A. performed the bioinformatic analysis of WGS data and developed TaqMan CNV PCR assays, S.F. developed and carried out the TaqMan CNV PCR experiments, G.S. performed some bioinformatic analysis of WGS data, F.P. conceived and supervised the study, R.S. conceived and supervised the bioinformatic analysis, wrote the manuscript. S.G. and L.P wrote the manuscript, conceived and supervised the entire study. All authors read and approved the final manuscript.

---

**Supplemental Figure S1.**
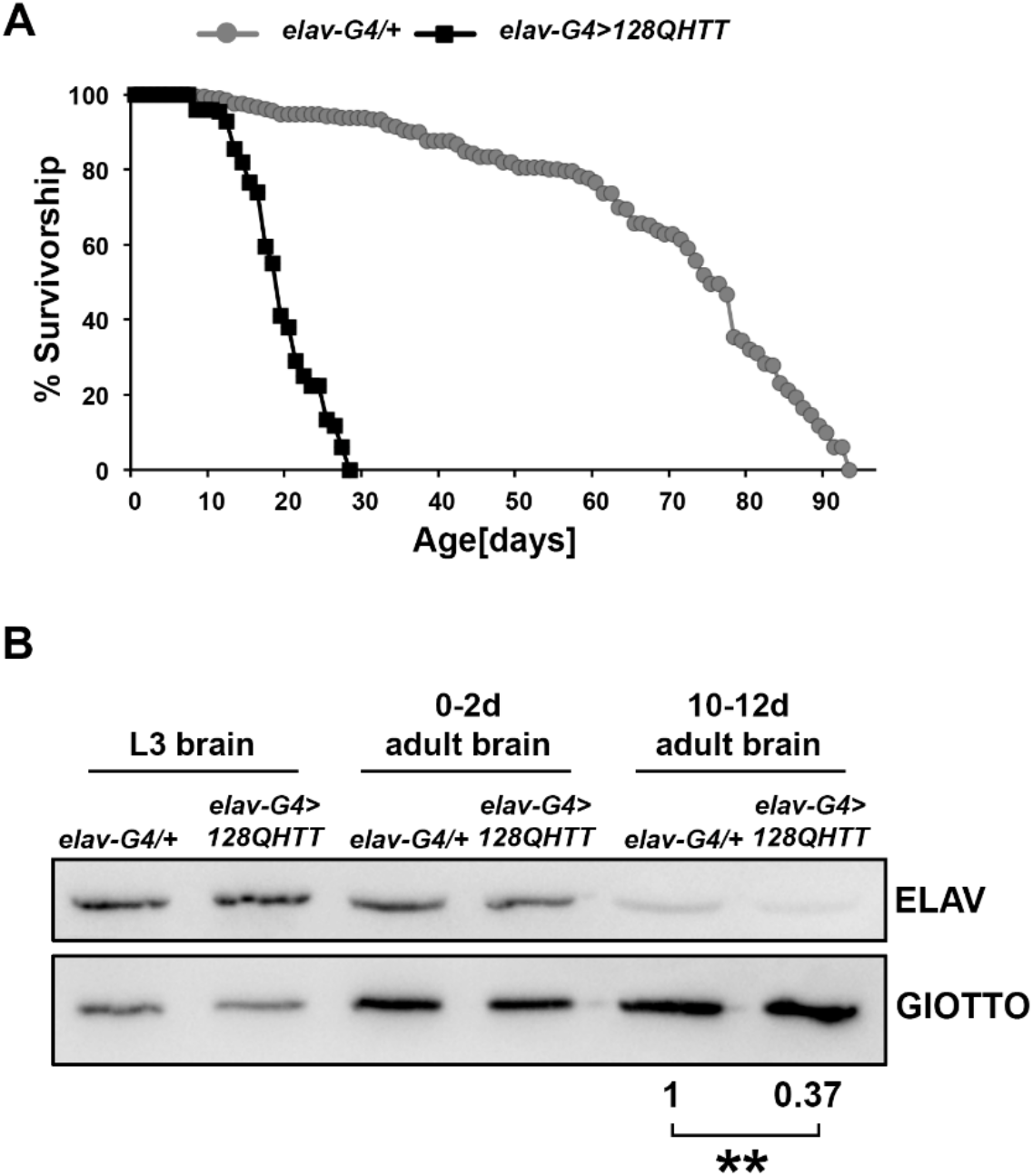
The expression of 128QHtt in neuronal cells reduces lifespan and results in cell death. (A) Survival curves of *elav-Gal4>128QHtt* flies (black squares) or *elav-Gal4/+* (grey circles). Data were from two independent experiments using at least 100 flies for each experiment. (***p < 0.001, Log-rank test). (B) Western blot analysis of ELAV protein expression in HD larval and adult brains. GIOTTO protein was used as a loading control. The result was expressed as means for at least three independent biological replicates (**p < 0.01, One-sample t-test).

**Supplemental Figure S2.**
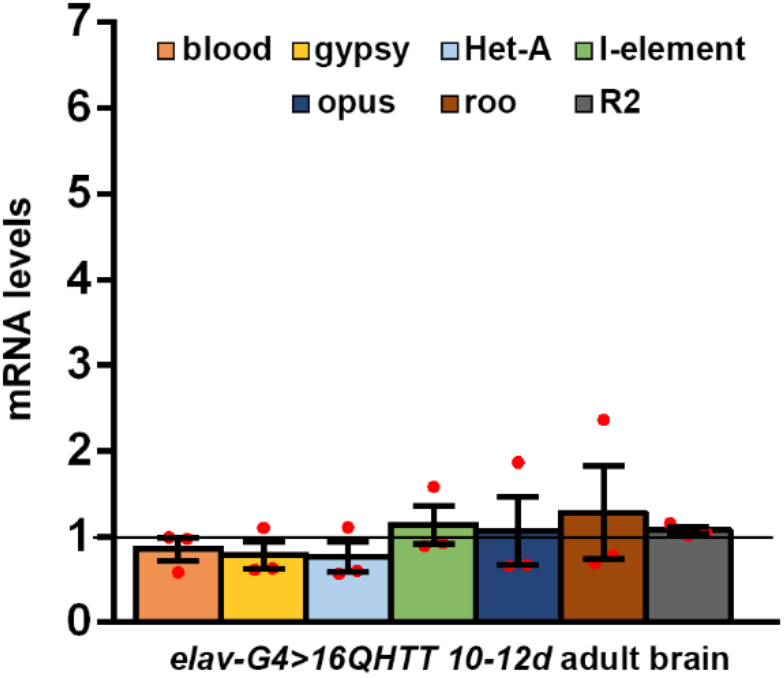
Neuronal expression of 16QHtt does not affect transposable element expression. Adult brains were analysed at 10-12d; transcript levels were normalized to *rp49* and displayed as fold change relative to *elav-G4/+* flies. Bar graph represents the mean ± SEM from at least three independent experiments (ns p>0.05, Unpaired t-tests). Red dots indicate individual data points. The black horizontal line indicates the Fold Change control value, set to 1.

**Supplemental Figure S3.**
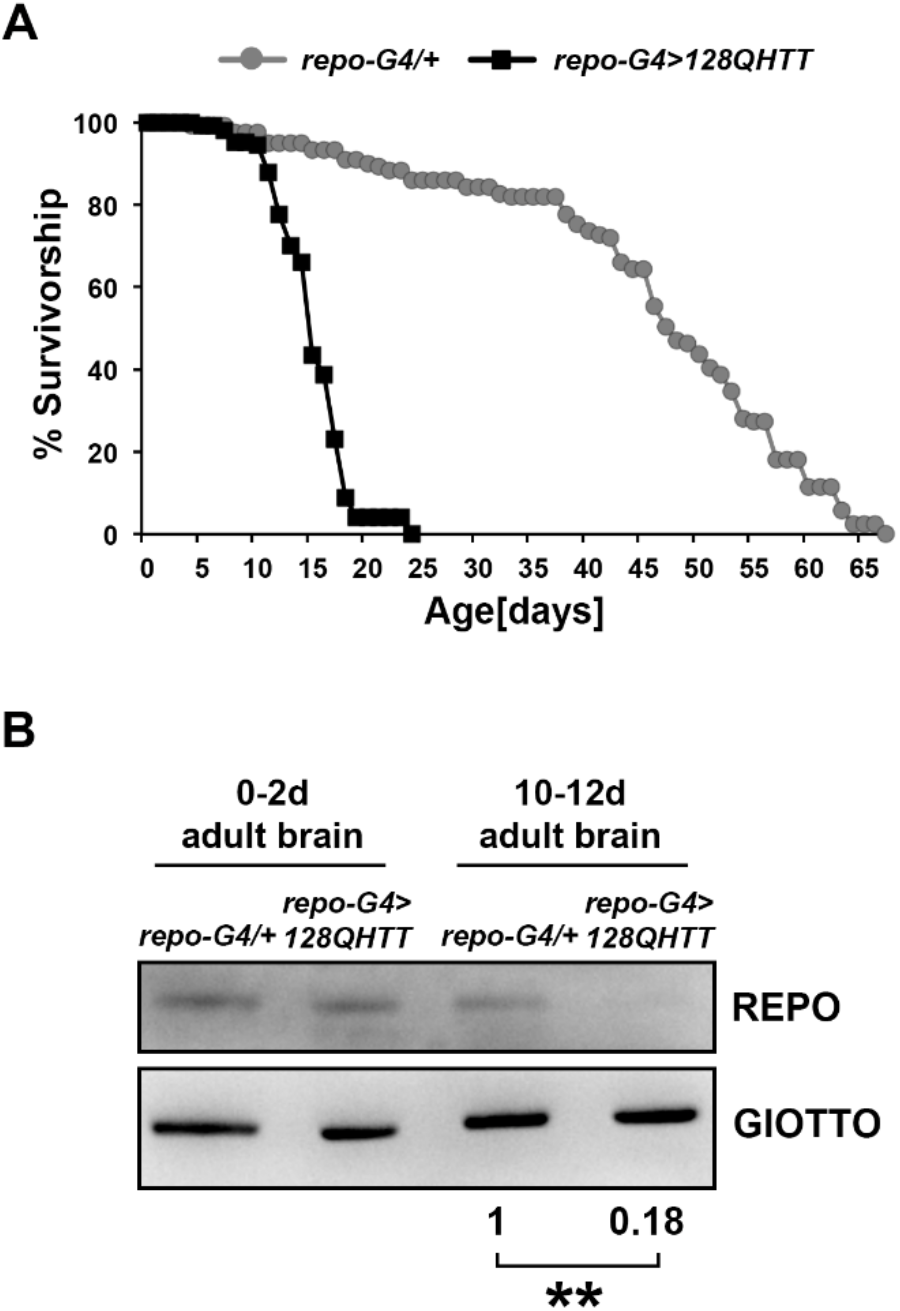
The expression of 128QHtt in glial cells reduces lifespan and results in cell death. (A) Survival curves of *repo-Gal4>128QHtt* flies (black squares) or *repo-Gal4/+* (grey circles). Data were from two independent experiments using at least 70 flies for each experiment. (***p < 0.001, Log-rank test). (B) Western blot analysis of REPO protein expression in HD larval and adult heads. GIOTTO protein was used as a loading control. The result was expressed as means for at least three independent biological replicates (**p < 0.01, One-sample t-test).

**Supplemental Figure S4.**
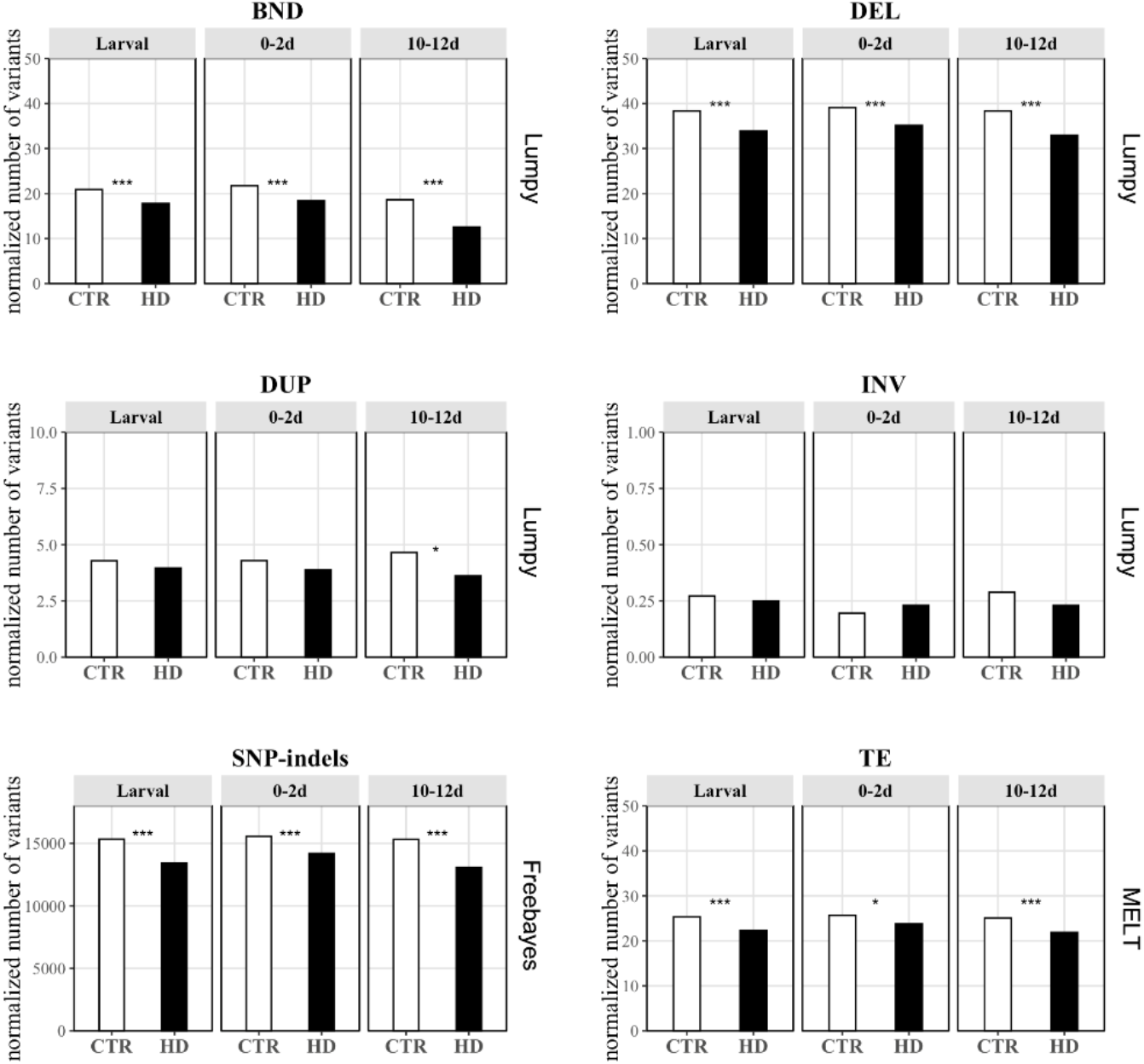
Number of structural variants. Break ends (BND), deletions (DEL), duplications (DUP), inversions (INV), SNP-indels and TE non reference ISs have been detected in each of the 6 samples analysed. A higher number of all the type of variants investigated have been detected in offspring CTR (white) than offspring HD (black) except for INV at 0-2d stage. The raw number of variants has been normalized on the total number of sequenced reads and multiplied by 1,000,000. Control samples are shown in white and HD in black. (* FDR < 0.05, ** FDR< 0.01, *** FDR< 0.001, Test of given proportion with Benjamin-Hochberg FDR correction).

**Supplemental Figure S5.**
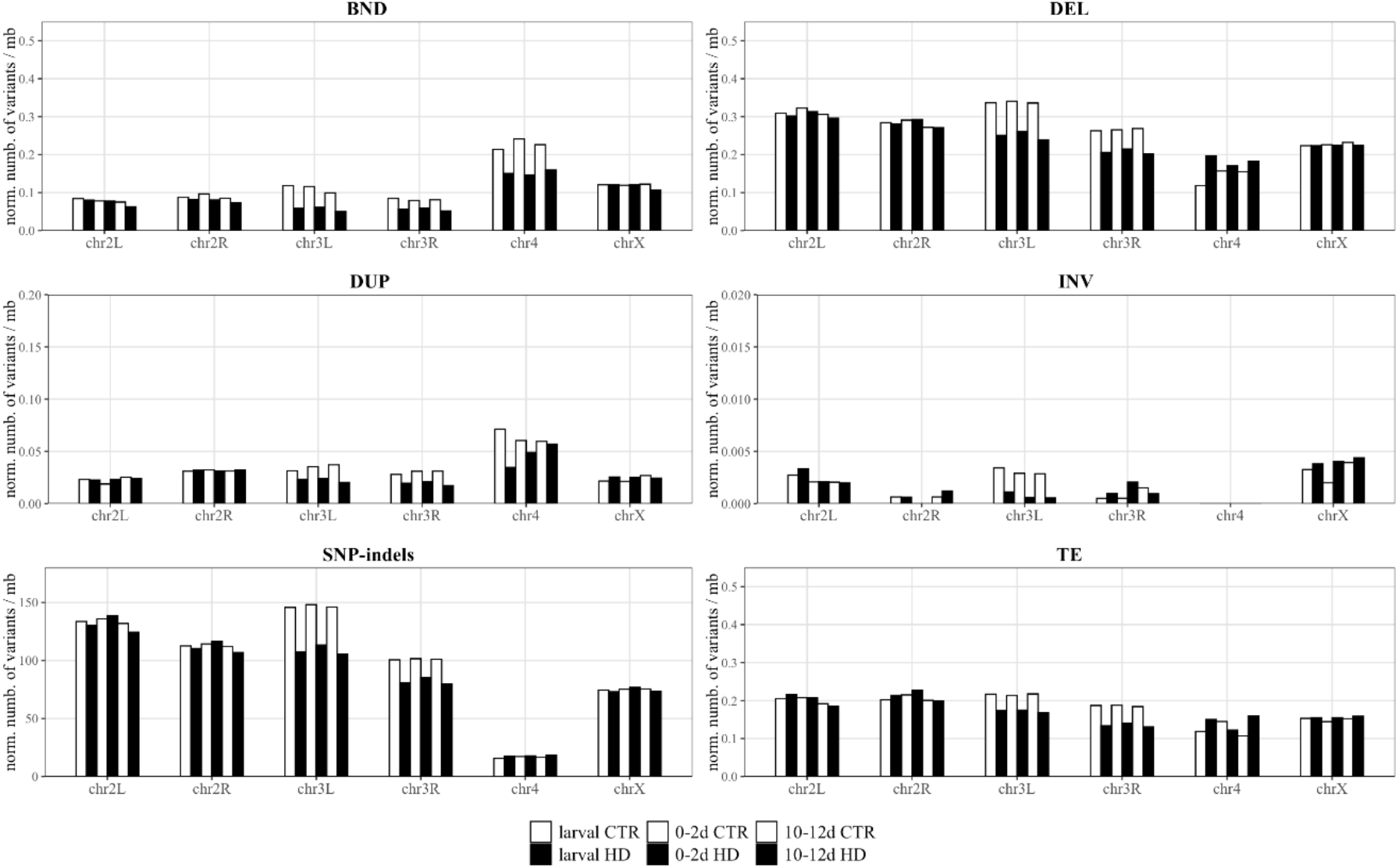
Number of structural variants on each *Drosophila* chromosome. All the variants investigated are enriched on chr3L and/or chr3R in offspring CTR samples (white). Offspring CTR flies carry a third balancer chromosome containing multiple inversions to avoid recombination. However, the balancer presence may have misled the variant call by the bioinformatic softwares here used. Raw number of variants has been normalized on total number of sequenced reads (multiplying by 1,000,000) and dividing by the chromosome length in megabases.

**Supplemental Figure S6.**
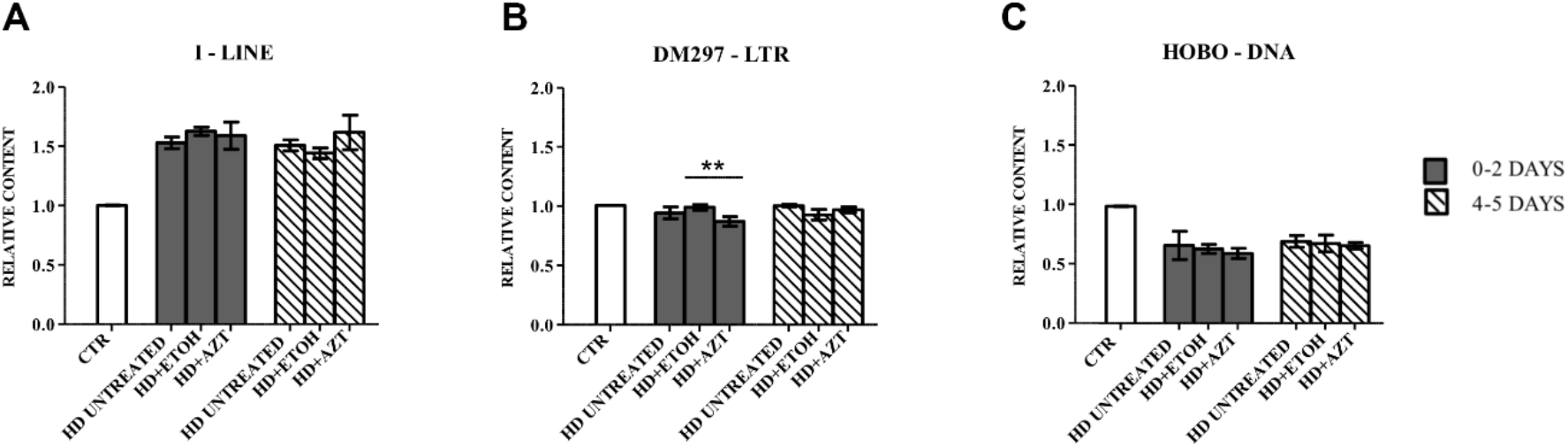
TaqMan CNV assay for I-element (retrotransposon), DM297 (retrotransposon), and HOBO (DNA transposon) on AZT-treated samples. (A to C) Taqman CNV assays as an exploratory assessment of RT-dependent CNVs of arbitrarily selected TE classes. I-element (A) and HOBO (C) did not show a statistically different cell content upon AZT treatment at 0-2d nor at 4-5d. DM297 showed a lower genomic level in samples treated with AZT compared to both control and vehicle (ethanol) at 0-2 days (B). CTR – brain from 10-12 days old offspring flies (white), for each AZT treatment (0-2d – dark grey, 4-5d – stripped bars) are reported HD untreated, HD treated with EtOH (AZT solvent) and HD +AZT. Cell content reported as ΔΔCt using as reference gene DMRT1C and as calibrator offspring larval CTR sample. (* p < 0.05, ** p < 0.01, *** p < 0.001, One Way Anova with Bonferroni post hoc test).

